# Combined-stress priming alters physiology, transcriptional dynamics, and acute thermal resistance in *Acropora cervicornis* corals

**DOI:** 10.64898/2026.04.16.719009

**Authors:** Serena Hackerott, Harmony Martell, Javier Rodriguez-Casariego, Jose M. Eirin-Lopez

## Abstract

Enhanced resilience conferred through sublethal stress pre-exposure may be crucial for reef building corals to cope with variable environments. The effect of stress priming on *Acropora cervicornis* thermotolerance was evaluated in the context of elevated temperature and ammonium enrichment, 3 ℃ and/or 6 μM above ambient, respectively. Primed corals were pre-exposed to each stressor individually or in combination for eight days, while non-primed corals remained at ambient conditions. After an eight-day recovery, primed corals and a subset of non-primed corals (naive) were subjected to an acute 15-hour thermal challenge. Coral metabolism, symbiosis, and gene expression were characterized throughout the experiment. Thermal tolerance was quantified as algal symbiont, chlorophyll, and live tissue retention, along with survival probability following acute heating. Primed corals were more likely to retain symbionts and chlorophyll after heat stress and also exhibited slower tissue loss. Moreover, thermal pre-exposure reduced the risk of tissue loss or predicted mortality. Apoptotic regulation differed between primed and naive corals during the initial and secondary heat exposures. Additionally, primed corals exhibited patterns of transcriptional resilience under acute thermal stress. Altogether, results provide support for the capacity of *A. cervicornis* to gain resilience through pre-exposure to ecologically relevant conditions as well as insights into the molecular mechanisms underpinning this process.

## Introduction

Stress priming (also referred to as pre-conditioning, pre-exposure, or stress-hardening) refers to sublethal environmental stress exposure that aims to increase an organism’s resistance to a subsequent stress event ^1^. The success of this intervention is predicated on the phenomenon of stress memory, where priming elicits a cellular response that is maintained and ultimately facilitates higher tolerance ^2^. Stress memory has been best described in terrestrial plants, and hardening techniques have been primarily used in agriculture ^3,4^. However, reef-building corals have also demonstrated a capacity for stress memory, likely through the same adaptive requirement as terrestrial plants: the need to rapidly acclimatize within variable environments without relocating ^5^. Further, given coral populations’ global decline and vulnerability to continued climate change, assisted evolution strategies, including stress hardening, have recently gained interest ^6–8^. However, many aspects of coral stress priming are not sufficiently understood to guide the application of this conservation intervention.

Coral stress memory was first observed in *Coelastrea aspera* corals, where elevated light intensity conferred higher thermal tolerance ^9^, which was maintained for 10 years ^10^. Such instances of cross-priming (also known as *trans*-priming), where the priming stressor is of a different origin than the secondary stressor, typically occur when both stressors trigger, and may be mitigated by, similar cellular pathways ^11^. The enhanced thermal tolerance in *C. aspera* colonies was associated with higher baseline levels of antioxidant enzymes as well as a stronger antioxidant and stress resistance (*e.g.,* heat shock proteins) response upon exposure to thermal stress ^12^. The combination of this prepared state and enhanced response resulted in less oxidative stress and protein degradation, and ultimately, less dysbiosis visible as coral bleaching. Oxidative stress mitigation is critical in determining the likelihood of coral bleaching, or the disruption of the symbiosis between the coral host and its symbiotic algae ^13^.

Since the initial observation, coral stress memory has been recorded across recurrent marine heatwaves ^14–17^ and coral thermal tolerance has been enhanced under controlled experimental conditions by pre-exposure to sublethal temperatures (*i.e., cis*-priming) ^18–21^, elevated irradiance ^22,23^, and herbicide ^24^. Yet, few studies have evaluated stress priming under simultaneous stressors ^25^, despite the likelihood of multiple sources of environmental stress impacting coral reefs ^26^. For instance, nutrient enrichment along with elevated temperature is a particularly ecologically-relevant combination ^27^ that has not yet been assessed. As elevated nutrients, especially nitrogen, have the potential to modulate coral vulnerability to heat stress ^28^, it is critical to evaluate thermal stress priming within the context of nutrient enrichment.

The first objective of this study was to evaluate the influence of pre-exposure to thermal, nutrient, and simultaneous stress on *Acropora cervicornis* thermotolerance. While once a regionally dominant species ^29^, *A. cervicornis*, or the staghorn coral, is now critically endangered ^30^ following precipitous population declines resulting from local and global stressors ^31^. Thus, *A. cervicornis* is currently the most targeted species for coral restoration ^32^ and represents an important candidate for assessing the outcomes of pre-exposure to ecologically relevant doses and sources of environmental stress. Pre-exposure to temperatures approximately two degrees below the bleaching threshold was predicted to enhance resistance to heat stress ^19,33^. The influence of nitrogen pre-exposure was more difficult to predict and could potentially reduce thermal tolerance ^34^, mitigate the negative effects of heat stress ^35^, or have no detectable effect on the response to elevated temperature ^36,37^. Similarly, the predicted benefit of increased thermotolerance conferred through thermal priming could be minimized, enhanced, or remain unaltered by the addition of elevated ammonium during stress priming.

The second aim of the study was to elucidate the molecular underpinnings of stress priming in corals. Three specific hypotheses were tested. The first was that if each type of stress pre-exposure enhanced thermal tolerance, this would be achieved through the activation of genes involved in similar biological processes across stressors ^11^. For example, both acute heat and ammonium stress induced caspase expression in *Pocillopora damicornis* corals ^38^, which may result in a similar prepared or “primed” state between stressors. The second hypothesis was that a portion of the transcriptomic response triggered by the priming exposure would be constitutively maintained over time as transcriptional preparation (*i.e.,* “frontloading”) ^39^ which could limit physiological stress occurring during the subsequent heat exposure ^5^. A potentially contrasting alternative hypothesis was that primed corals would instead mount a faster and/or stronger response at the onset of the second heat exposure, in turn, reducing the resulting physiological stress ^1^. Lastly, the third hypothesis was that the enhanced thermal tolerance of the “primed” state would be associated with transcriptional resilience, or a faster return to baseline levels of gene expression following stress exposure. Such transcriptional resilience has also been associated with stress resistance in corals, likely indicating a faster re-establishment of homeostasis ^40^.

## Results

### Environmental Stress Priming

Ramets of three A. cervicornis genets (**Text S1**) were distributed across four potential treatments: ambient (control), elevated temperature, elevated nutrients, or the combination of both (**Figure S1**). Priming treatments were maintained for eight days (**Text S2, Figures S2** and **S3**) and then returned to ambient for an eight-day recovery period. During the priming phase, the cumulative heat exposure within the elevated temperature tanks (temperature- and combined-primed) was 17.222 ± 0.277 (□ * day) (**Table S1**), and the water temperature profiles were significantly different from those of the ambient tanks (**Figure S3, Table S2**). Additionally, profiles of ammonium concentration in the tanks pre-exposed to elevated nutrients (nutrient-and combined-primed) were significantly different from those of the ambient tanks. The maximum concentration of ammonium was approximately 2.7 μM lower than the intended exposure (**Table S1**), which may be due to a higher rate of nutrient uptake than expected ^34^.

### Priming Influence on Physiology

Chlorophyll concentration was reduced following stress priming, and this reduction persisted following the recovery phase (**Figure S4**, **Table S3**). Evidence of sublethal stress was also detected in coral metabolism following priming. The ratio of gross photosynthesis to respiration (P_Gross_:R) increased in all primed treatments relative to the ambient control, driven by lower respiration rates in primed corals (**Figure 1, Table S4**). Such a pattern was still apparent in the nutrient-primed treatment after five days of recovery. Yet, no significant differences in P_Gross_:R or respiration were detected following the full (eight-day) recovery phase (**Figure S4**). Additionally, gross photosynthesis was not influenced by the priming treatments at any time point (**Figures 1, S4, Table S4**).

**Figure 1.**
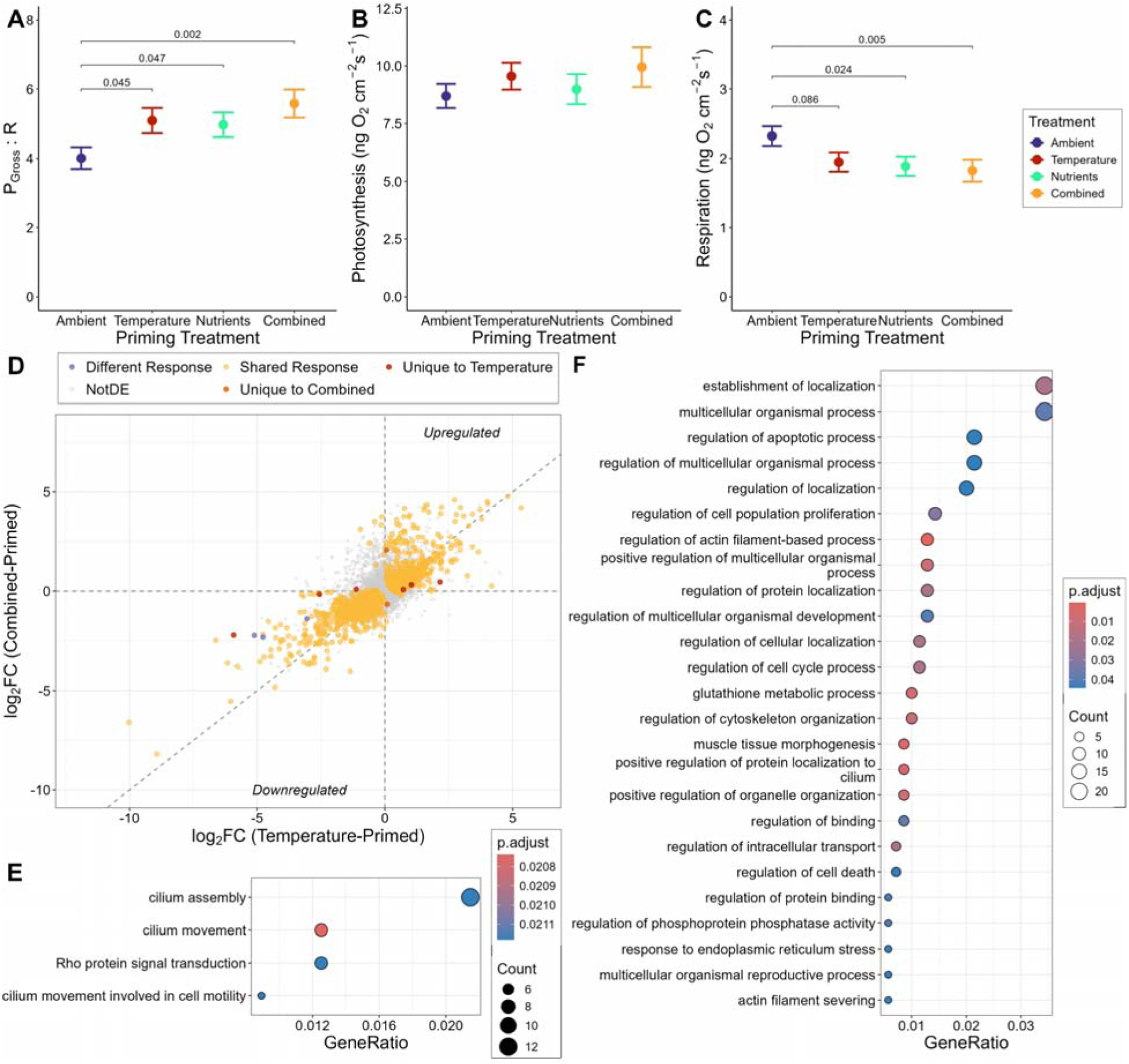
Coral metabolism and gene expression after stress priming. **A)** The ratio of gross photosynthesis to respiration (P_Gross_ : R), **B)** Gross photosynthesis (ng O_2_/cm^2^/s), and **C)** respiration (ng O_2_/cm^2^/s) for each treatment at the post-priming time point. Points represent averages, and error bars represent the standard error, each colored by priming treatment. For cases when the response metric was significantly influenced by the priming treatment, p-values comparing each primed treatment to the ambient control are shown (**Table S4**). **D)** Gene expression (log_2_ fold change [log_2_ FC]) of the temperature-primed corals (x-axis) versus the combined-primed corals (y-axis), relative to the ambient control corals. Points are colored by the uniqueness of the response between the treatments being compared (grey: genes not differentially expressed, yellow: genes differentially expressed in the same direction in both treatments, purple: genes differentially expressed in both treatments but in different directions, red: genes differentially expressed only in the temperature-primed treatment, and orange: genes differentially expressed only in the combined-primed treatment). **E-F)** Gene ontology (GO) terms of biological processes enriched (**Table S6**) in the shared transcriptomic response to priming in both the temperature- and combined-primed treatments, including upregulated (E) and downregulated (F) genes. For each term, the dot color corresponds to the adjusted p-value, and the size denotes the number of significant genes within the term.

The observed differences in coral symbiosis and metabolism were not due to changes in symbiont density (**Figure S4**, **Table S3**) or shifts in symbiont community composition following stress priming (**Table S5**). All samples were dominated by *Symbidinium fitti* (A3), with some sequence variants present in low relative abundance across samples (**Figure S5**). At the level of sequence variants, symbiont communities differed between the three genotypes (**Table S5**). There was also an effect of time point, yet this cannot be distinguished from potential intra-colony variation across different ramets sampled through time. However, the symbiont community described at the level of ITS2 profiles did not differ between priming treatments, genotypes, or time points (**Figure S5, Table S5**).

Pre-exposure to elevated water temperature and ammonium induced transcriptomic responses in primed corals. After the priming phase, 1,407 genes (out of the 16,191 genes retained for analysis) were differentially expressed (627 up- and 780 down-regulated) in the temperature-primed corals relative to the ambient corals. A total of 865 genes were differentially expressed (396 up- and 469 down-regulated) in combined-primed corals. Only 15 differentially expressed genes (DEGs) were detected in the nutrient-primed corals (all downregulated). The response to stress priming was largely shared between the two treatments pre-exposed to elevated temperature, with only 11 genes differing in response between the temperature- and combined-primed treatments (**Figure 1D**). The shared response to priming included genes associated with movement and localization, regulation of apoptosis and cell proliferation, as well as oxidative and endoplasmic reticulum stress (**Figure 1E-F, Table S6**). Following recovery, only 1 gene remained significantly differentially expressed in each of the primed treatments, therefore gene expression at this time point was not further analyzed.

### Thermotolerance After Priming

Following recovery, primed corals and a subset of previously ambient corals (naive) were challenged with acute heat stress to evaluate changes in thermal tolerance (**Figures S1** and **S3**). During the acute thermal challenge, the cumulative heat exposure was 3.251 ± 0.345 (□* day) (**Table S1**) and water temperature profiles were significantly different in the heated tanks compared to the control tanks (**Figure S3, Table S2**). By the end of the thermal challenge, primed corals were 2.94-6.56 times more likely to have retained 100% symbiont density (i.e., control levels) compared to the naive corals, and 4.16-6.61 times more likely to retain 100% chlorophyll concentration (**Table 1 and S7**). Corals in the combined-primed treatment retained significantly more symbionts and chlorophyll than naive corals and the temperature- and nutrient-primed corals had marginally higher odds (p < 0.1) of retaining symbionts and chlorophyll, respectively (**Figure 2, Table S7**).

**Figure 2.**
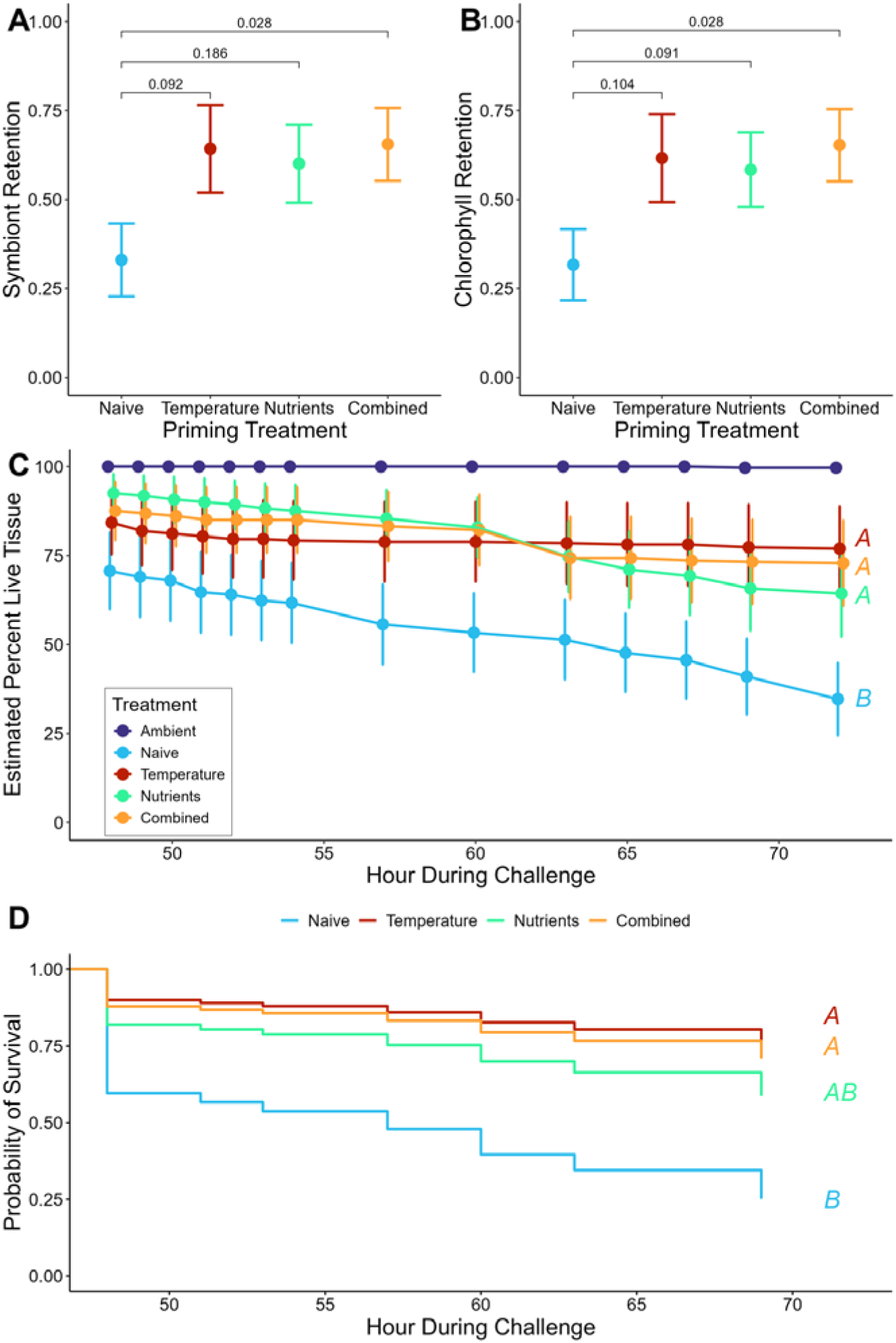
Retention of symbionts, chlorophyll, and live tissue. Proportion retained ((Treatment / Control); see **Figure S6** for measured values) of **A)** symbiont density (cells/cm^2^) and **B)** chlorophyll concentration (μg/cm^2^) at the post-challenge time point. P-values (**Table S7**) comparing each primed treatment to the naive treatment are shown. For panels **A-C**, points represent averages and error bars represent the standard error, each colored by priming treatment. **C)** Estimated live tissue (percent of fragment surface area) throughout hours 48-72 of the thermal challenge phase. Capital letters indicate significant pairwise differences in the rate of tissue loss compared between treatments that were heated during the thermal challenge (**Table S7**). **D)** Survivorship curves (cox proportional hazard) of the probability of survival throughout the thermal challenge. Curves are colored by priming treatment. Capital letters indicate significant pairwise differences in hazards between treatments (**Table S7**).

**Table 1.**
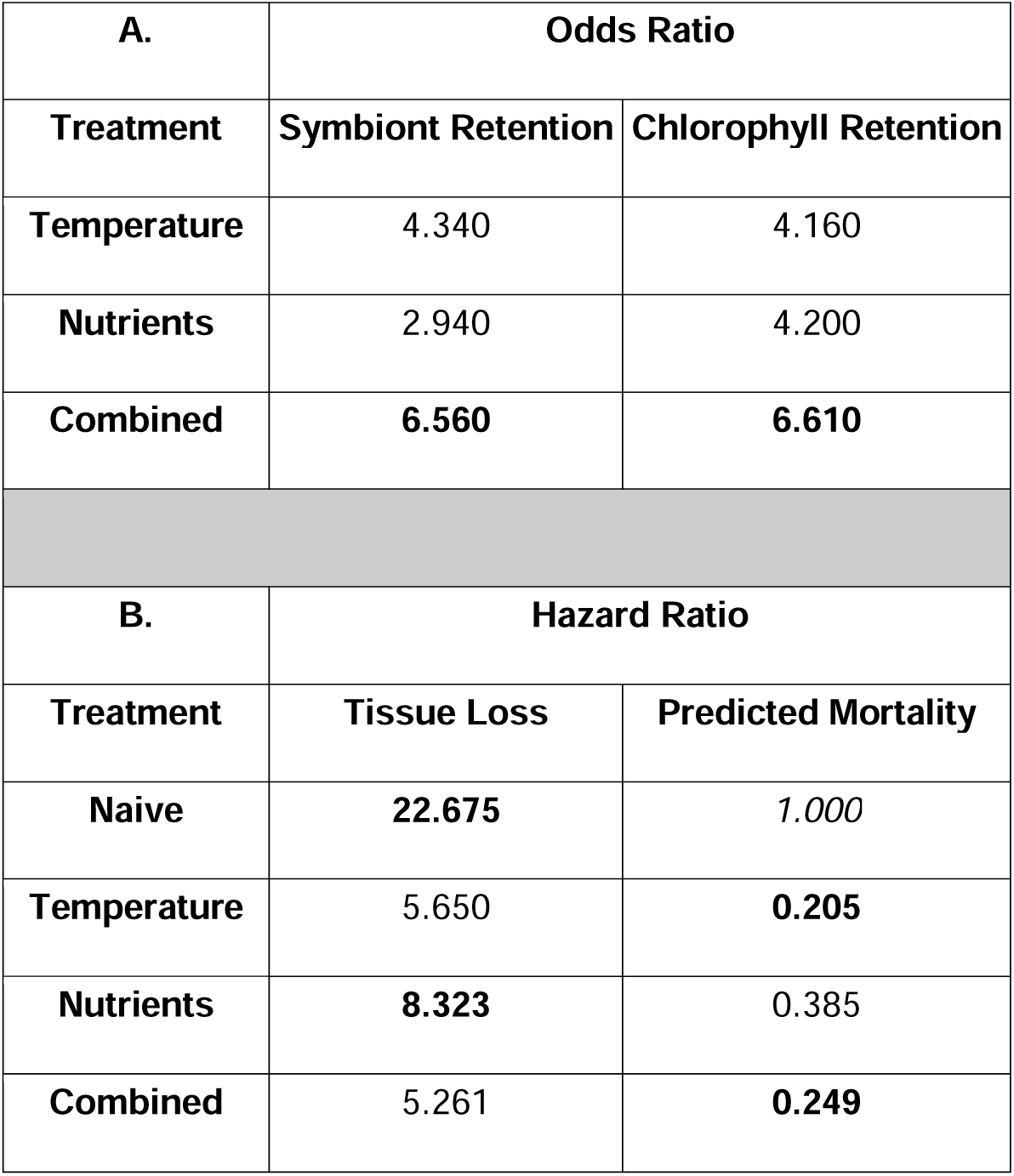
Odds and hazard ratios of bleaching, tissue loss, and predicted mortality. **A)** The odds ratio of each primed treatment retaining symbionts (**left**) and chlorophyll (**right**) after the thermal challenge, relative to the odds of retention in the naive treatment. Ratios were determined from generalized linear mixed effects models (**Table S7**). **B)** The hazard ratios of experiencing tissue loss (**left**) and predicted mortality (tissue loss > 20%) (**right**) throughout the thermal challenge. For the **left** panel, the relative risk of tissue loss in each of the heated treatments (*i.e.,* naive and primed treatments) is compared to the risk in the ambient (control) treatment. For the **right** panel, the risk of predicted mortality in each primed treatment, relative to the risk in the naive treatment (*ratio relative to risk of 1*), is shown. Ratios were determined from cox proportional hazards models (**Table S7**). For panels **A** and **B**, odds and hazards significantly different from the reference treatments are in bold.

Throughout the thermal challenge, tissue loss progressed faster in the naive corals compared to corals in each of the primed treatments (**Figure 2**, **Table S7**). Indeed, naive corals were 22.68 times more likely to experience tissue loss compared to corals in the control treatment (**Table 1**). Temperature- and combined-primed corals were less likely to experience tissue loss compared to naive corals, and the risk of tissue loss in these primed corals was actually not significantly different than that of ambient control corals (**Figure S6**, **Table S7**). Furthermore, primed corals in these elevated temperature treatments were significantly more likely to survive than the naive corals, with less than a quarter of the risk of mortality (**Figure 2**, **Table 1**). The results of nutrient priming on thermotolerance were intermediate. Nutrient-primed corals were significantly more likely to experience tissue loss compared to the ambient corals (**Table 1**), similar to the naive corals. Yet, the risk of tissue loss and predicted mortality in the nutrient-primed corals did not differ from that of corals in the other two primed treatments, nor from the risk of the naive corals (**Figures 2 and S6, Table S7**).

As temperatures returned to ambient levels following the acute heat stress (thermal challenge time point C6), all heated corals displayed large transcriptional responses. Relative to the ambient control, 7,259 genes were differentially expressed (2,898 up- and 4,361 down-regulated) in the naive treatment (out of the 15,584 genes retained for analysis). Similarly, 7,440 genes were differentially expressed (3,078 up- and 4,362 down-regulated) in the nutrient-primed treatment. Corals pre-exposed to elevated temperature displayed comparatively fewer DEGs (temperature-primed: 6,252 [2,570 up- and 3,682 down-regulated] and combined-primed: 7,016 [2,799 up- and 4,217 down-regulated]). While the majority of DEGs were shared between naive and primed corals, stress priming did alter the response to acute heat exposure. The temperature-primed response was comparatively more muted than the naive response, with only 8 genes unique to and/or more strongly expressed in that treatment compared to the naive treatment, where 114 DEGs were uniquely and/or relatively more strongly expressed (**Figure 3A**). The nutrient-primed response differed from the naive response in more DEGs, with 205 and 148 uniquely and/or more strongly expressed in the nutrient-primed and naive treatments, respectively (**Figure 3B**). Interestingly, by the end of the acute heat exposure, the combined-primed response was unique in only 10 genes compared to the naive response, and only 4 genes were unique or relatively more responsive in the naive treatment (**Figure 3C**).

**Figure 3.**
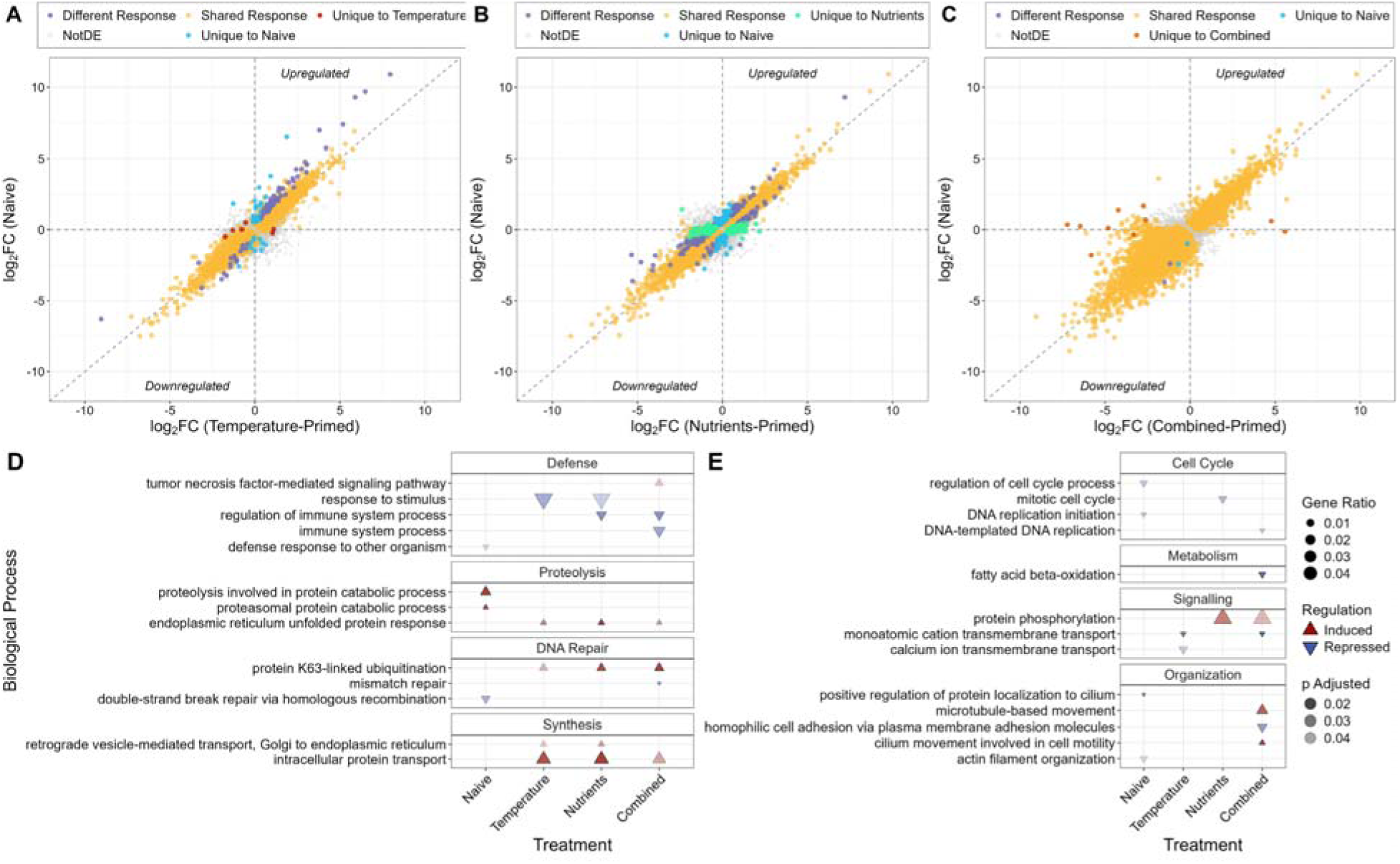
Transcriptomic responses to acute heat stress. **A-C)** Gene expression (log_2_ fold change [log_2_ FC]) of the naive corals (x-axis) versus the primed corals (y-axis), relative to the ambient control corals. Primed treatments are across columns (**A**: temperature-, **B**: nutrients-, **C**: combined-primed). Points are colored by the uniqueness of the response between the treatments being compared. **D-E)** Gene ontology (GO) terms of biological processes enriched at the end of the thermal challenge, which differed between naive and primed treatments (see **Figure S7** and **Table S8** for full enrichment results). Induced terms are shown in upward red triangles, and repressed terms are shown in downward blue triangles. The triangle size corresponds to the ratio of significant genes within the term, and the transparency corresponds to the adjusted p-value. GO terms are organized by similarity.

Biological processes involved in the core heat stress response, common across heated treatments, included those related to protein synthesis and degradation, cell cycle, and signalling (**Figure S7, Table S8**). However, there were also notable differences in the primed vs naive response to acute heat stress, particularly in defense and DNA repair functions (**Figure 3D-E**). For example, genes related to “defense response to another organism” were only enriched in the naive treatment, while genes related to immune system processes and response to stimulus were enriched in at least two of the three primed treatments. Additionally, all primed treatments upregulated genes relative to the endoplasmic reticulum unfolded protein response, protein K63-linked ubiquitination, and intracellular protein transport, which were not enriched in the naive treatment.

### Primed Heat Stress Response and Dynamics

Throughout the thermal challenge, the transcriptomic response of the heated corals peaked at thermal challenge time point C4, which corresponded to the peak heat stress (**Table 2**). Yet, differences between temperature-primed and naive corals were apparent and peaked earlier during the acute heat exposure (**Figure S8**). For example, direct differential expression comparisons between primed and naive corals detected 61 and 222 DEG at time points C2 and C3, respectively. Moreover, 28 and 50 genes were uniquely and/or more strongly expressed in the primed corals at C2 and C3, respectively, in addition to 19 and 92 uniquely and/or more strongly expressed genes in naive corals at these time points. Notably, genes related to the regulation of apoptosis were only enriched in the primed treatment at time point C2 and remained more strongly enriched at C3 relative to the naive corals (**Figure S9, Table S9**). Additional early heat stress responses that were unique to or stronger in temperature-primed corals included protein ubiquitination, cell cycle regulation, and membrane and cellular organization.

**Table 2.**
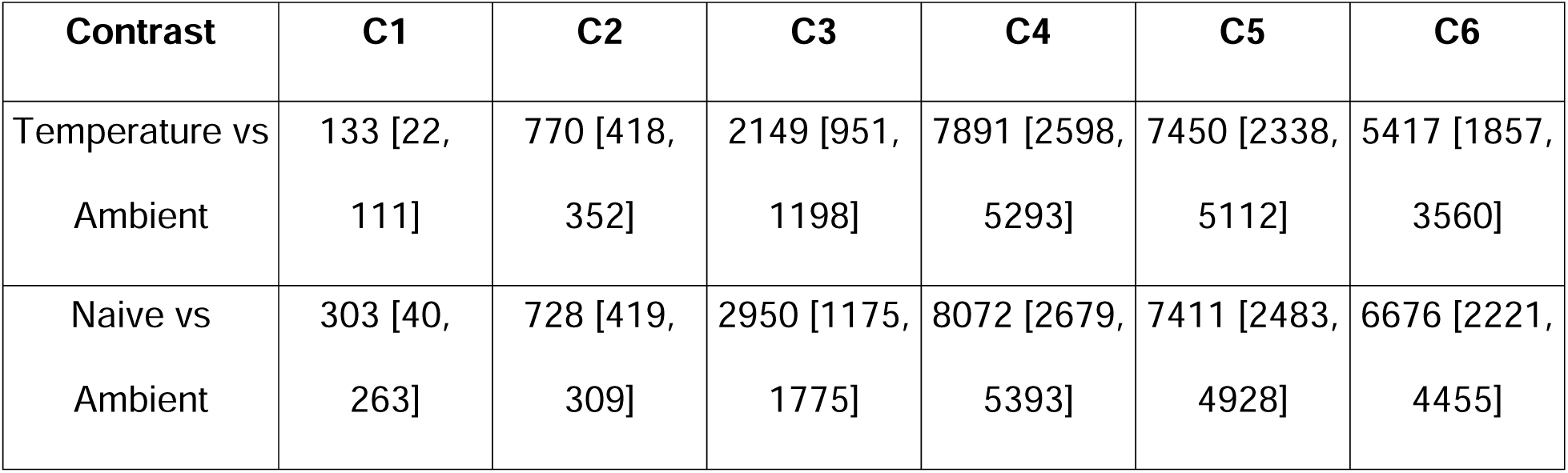
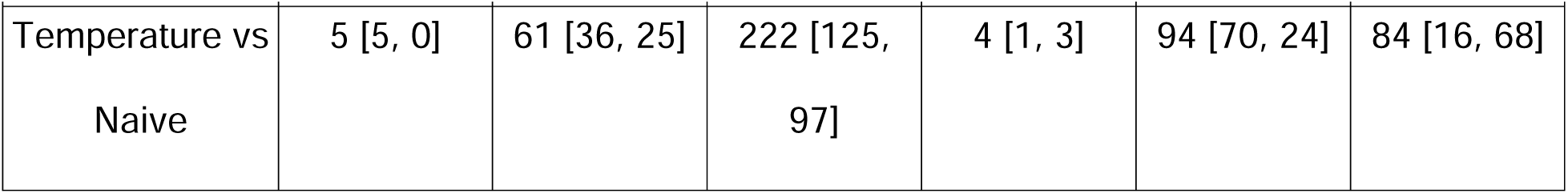
Differential gene expression throughout thermal challenge. Number of significantly differentially expressed genes (DEG) throughout the thermal challenge (time points C1-C6) across three contrasts of interest. Values are shown as total DEG [upregulated, downregulated] where regulation direction is in terms of the first treatment contrasted against the second.

| Contrast | C1 | C2 | C3 | C4 | C5 | C6 |
| --- | --- | --- | --- | --- | --- | --- |
| Temperature vs Ambient | 133 [22, 111] | 770 [418, 352] | 2149 [951, 1198] | 7891 [2598, 5293] | 7450 [2338, 5112] | 5417 [1857, 3560] |
| Naive vs Ambient | 303 [40, 263] | 728 [419, 309] | 2950 [1175, 1775] | 8072 [2679, 5393] | 7411 [2483, 4928] | 6676 [2221, 4455] |
| Temperature vs Naive | 5 [5, 0] | 61 [36, 25] | 222 [125, 97] | 4 [1, 3] | 94 [70, 24] | 84 [16, 68] |

Later during the thermal challenge, as water temperatures decreased towards the ambient baseline (time points C5 and C6, respectively), 94 and 84 genes directly differed between the primed and naive treatments (**Table 2**). These differences were largely driven by genes uniquely and/or more strongly expressed in the naive corals (C5: 52 genes and C6: 77 genes) relative to the primed-response (**Figure S8**). Further, expression within gene modules that were more highly correlated with the naive treatment compared to the primed treatment (**Figure S10**) tended to diverge later during the challenge (**Figure 4A**). Four of these key gene modules showed evidence of transcriptional resilience in primed corals, including three modules of upregulated genes (2,278 genes total) and one large module of 5,068 of downregulated genes. Genes within the upregulated modules were associated with transcription, proteolysis, defense, DNA repair, as well as cell cycling, signaling, and organization (**Figure 4B**). Genes in the downregulated module were largely associated with metabolism, DNA repair, and the cell cycle (**Figure 4C**).

**Figure 4:**
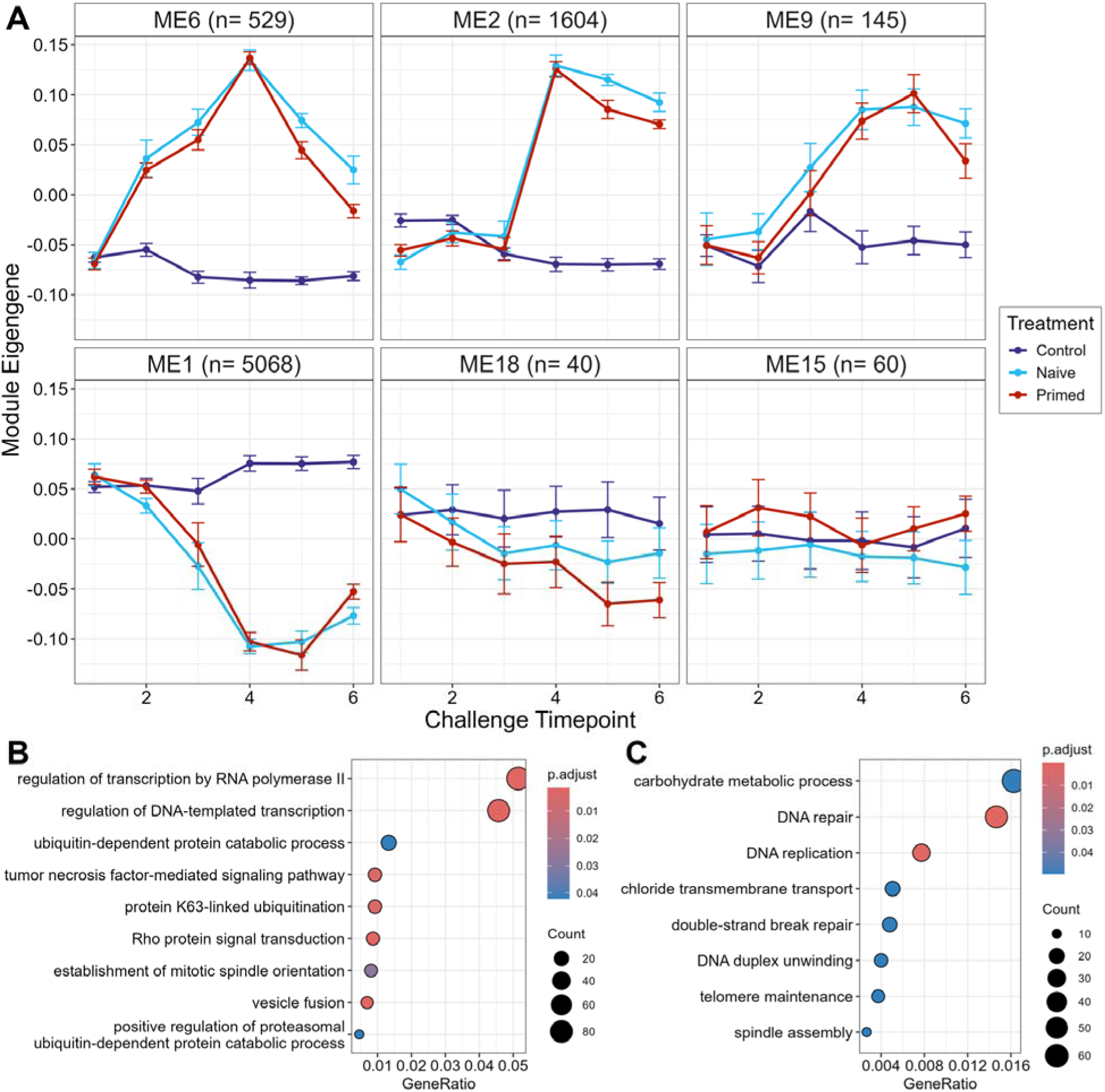
Expression dynamics and functional enrichment throughout thermal challenge. **A)** Modules of genes displaying similar expression patterns throughout the thermal challenge, identified through weighted gene coexpression network analysis (WGCNA), where expression levels (module eigengenes) of gene groups differed in strength (Naive > Primed: ME6, 2, 9, 1; Primed > Naive: ME18) or direction (ME15) between primed and naive treatments (see **Figure S10** for full WGCNA trait association results). The number of genes per module is noted. Gene ontology (GO) terms of biological processes enriched in modules where expression was stronger in the naive treatment compared to the temperature-primed in **B)** upregulated (ME6, 2, 9) and **C)** downregulated (ME1) gene groups (**Table S10**).

## Discussion

Pre-exposure to temperatures approximately two degrees below the bleaching threshold ^19,33^ significantly increased *Acropora cervicornis* acute thermal tolerance, as predicted. Corals primed with elevated temperature were approximately 4-5 times less likely to bleach or exhibit tissue loss during the thermal challenge, compared to corals that were not pre-exposed to stress (*i.e.,* naive). Furthermore, temperature stress priming significantly reduced the predicted heat-induced mortality risk. Two other recent studies have assessed the effect of thermal priming and acclimation (*i.e.,* pre-exposure with no recovery period) on *A. cervicornis* thermotolerance. Long-term (89 days) acclimation to variable temperatures (28-31 □) delayed bleaching during a subsequent heat stress assay ^41^. In contrast, bleaching occurred earlier in corals acclimated to a static elevated temperature (28 □) for the same timeframe. However, priming exposures of similar temperatures (29 or 31 □) for much shorter durations (1 or 2 days) did not significantly reduce bleaching during an acute thermal challenge ^42^. Together, these and our results highlight the importance of the pre-exposure dose (*e.g.*, magnitude, duration, and frequency) on the outcome of stress priming ^5^, but the optimal dose has yet to be established. Still, the reduced risk of bleaching and mortality observed in these medium-term and previous long-term studies provides promising support for the potential to enhance resilience in this critically endangered species through thermal priming.

Following thermal priming, genes related to localization and movement, regulation of cell death and proliferation, and oxidative and endoplasmic reticulum stress were differentially expressed in pre-exposed corals. Apoptosis plays a critical role in the cellular pathway that results in coral bleaching ^43^, and the regulation of this process has been linked to heat stress outcomes in corals ^44–46^. Further, the relative expression of pro-survival to pro-death genes regulating apoptosis during heat exposure has been identified as a key driver of the elevated resilience conferred by thermal priming ^47,48^. By the end of the thermal challenge, primed corals differed from naive corals in the regulation of a few key processes, especially defense responses. Additionally, primed corals displayed a divergent endoplasmic reticulum (ER) unfolded protein response and also differentially regulated protein K63-linked ubiquitination. The ER unfolded protein response has been identified as a key line of defense in Pacific congener acroporid corals for coping with recurrent environmental stress ^49^. Additionally, K63□linked ubiquitination mediates non-proteolytic stress signaling and translational control during oxidative stress ^50^, a core feature of bleaching physiology ^51,52^. Together, these results elucidate the pathways altered by stress priming which likely facilitated the enhanced thermal resistance observed in this study.

Nutrient enrichment during thermal priming may have augmented bleaching resistance benefits. Corals pre-exposed to elevated temperature in concert with elevated ammonium (combined-primed) were significantly more likely to retain symbionts and chlorophyll concentrations during the bleaching assay (approx. 6.5 times more than naive corals). Yet, the risk of tissue loss and predicted mortality was comparable between corals pre-exposed to elevated temperature with or without nutrient enrichment. Elevated ammonium has the potential to ameliorate the detrimental effects of heat stress in corals through reducing oxidative stress ^35^ and promoting anti-inflammatory fatty acid biosynthesis ^53^. Such mechanisms may have contributed to the observed advantage in symbiont and chlorophyll retention under heat stress by reducing the physiological costs of heat pre-exposure. As coastal nutrient enrichment is common in developed locations ^54^, the potential net positive (or neutral) influence of moderately elevated ammonium during thermal priming is an important consideration for conducting *in-situ* coral stress hardening.

Pre-exposure to elevated nutrients at ambient temperatures may provide a carry-over benefit when corals are subsequently challenged with heat stress. Corals primed with elevated ammonium were approximately 3-4 times less likely to bleach compared to naive corals and tissue loss progressed more slowly in nutrient-primed compared to naive corals. Yet, the concentration, source, and relative abundance of critical nutrients, including nitrogen and phosphorus, all modulate the influence of nutrient enrichment on coral symbiosis, and ultimately, the vulnerability of coral holobionts to thermal stress ^28^. In fact, acclimation at a higher magnitude of ammonium (10 μM above ambient, compared to an increase of ∼3.3 μM in this study) for a longer duration (78 days, compared to 8 days in this study) resulted in decreased heat resistance in *A. cervicornis* corals ^34^. In contrast, long term (months-weeks) acclimation periods to low-level nutrient enrichment have increased thermal performance ^55^ and delayed bleaching ^56^ in multiple Pacific coral species. The influence of nutrient priming is also likely to vary between sources of nitrogen. For example, nitrate enrichment increased bleaching prevalence, while enrichment of urea did not alter bleaching severity ^37^. Additionally, the detrimental effect of nutrient enrichment on thermal tolerance more commonly occurs when nitrogen (especially nitrate) is elevated disproportionately to phosphorus ^57,58^. Such nutrient imbalances can cause phosphorus starvation, resulting in destabilization of photosynthetic machinery ^59^ along with negative feedbacks, including a reduced capacity to repair oxidative damage ^60^ or assimilate dissolved inorganic phosphorus ^61^. The influence of nutrient pre-exposure on coral thermal tolerance should therefore be further assessed under a broader range of environmentally-relevant nutrient concentrations.

This study also evaluated key hypotheses describing the transcriptional response of stress-primed corals, including that thermotolerant primed corals would display 1) transcriptional frontloading through constitutive gene regulation as the mechanism of perpetuating stress memory, or 2) a stronger initial response at the onset of recurrent stress. It was also predicted that each of these enhanced response mechanisms may be accompanied by 3) higher transcriptional resilience during stress exposure, with a faster return to baseline gene expression. We did not find evidence of transcriptional frontloading throughout this relatively short-term study, demonstrated by the lack of DEG’s detected in the primed treatments after 8 days of recovery. Similarly, acclimation to temperatures 3 □ above ambient for 7-11 days increased the resistance of *Acropora nana* corals to acute heat stress without transcriptional changes detectable prior to the thermal challenge ^62^. While constitutive gene expression has been implicated in the higher resilience of corals adapted and/or acclimated to variable stress regimes over longer periods ^39,63^, alternative cellular mechanisms were likely responsible for the transmission of stress memory observed in the present study. Reviewed in ^5^, these may include post-translational regulation leading to the persistence of stress-responsive proteins as well as changes in gene accessibility through epigenetic modifications, which both have the capacity to facilitate an enhanced defense to recurrent stress. In fact, post-translational histone modifications (PTM’s), changes to nucleosome composition, and poised polymerase can maintain “transcriptional memory” through stress priming in plants, without constitutive gene expression ^2,64,65^. Thus, epigenetic gene and chromatin regulation should be evaluated as putative mechanisms for perpetuating the stress-primed state in corals.

Thermotolerant primed corals in the present study mounted targeted early defense mechanisms near the onset of acute heat exposure. Specifically, genes related to apoptosis regulation were enriched sooner and more strongly in primed corals compared to naive corals within the first 4-7 hours of the acute thermal challenge. Similarly, stress pre-exposed *Pocillopora acuta* corals shifted towards the “pro-survival” regulation of apoptosis 3 hr into heat exposure ^47^. As early and fine-tuned regulation of apoptosis is a key predictor of coral heat stress outcomes ^66,67^, such rapid enrichment in primed corals likely contributed to the lessened heat-induced physiological stress observed in this study. Interestingly, this function was also one of the most enriched at the end of the initial priming phase, further supporting the hypothesis that pre-exposure to sublethal stress can establish transcriptional contexts which are more responsive upon detection of recurrent stress ^3^.

The enhanced bleaching resistance and improved survival outcomes in primed corals may have also been linked to transcriptional resilience observed towards the end of acute heating. As water temperatures began to decline, transcriptional differences between primed and naive corals consisted primarily of stronger expression remaining in the naive treatment. Genes that began to return more quickly to baseline levels in primed corals included those related to transcription, proteolysis, defense, and DNA repair. This pattern of resilience can be interpreted as higher efficiency of these functions in primed corals, requiring less persistent expression to reestablish homeostasis ^49^. Indeed, transcriptional resilience has been associated with coral thermal resistance ^68,69^, especially across recurrent stress events ^70^. Interestingly, genes involved in Rho protein signal transduction were enriched following thermal priming and also displayed patterns of transcriptional resilience during the subsequent acute heat stress. Rho GTPases regulate cytoskeletal remodeling, vesicle trafficking, and cell to cell junction stability, which are essential for maintaining tissue structure and coral host to symbiont cellular organization ^71^. Rho signaling therefore likely plays an important role in cellular triage under thermal stress ^43^. The observed transcriptional resilience during acute heat stress is consistent with a primed, hormetic state, where prior exposure reduced the need for sustained transcriptional activation and ultimately supported cellular stability and stress tolerance.

In summary, the present work demonstrated that pre-exposure to sublethal temperatures elevated the acute thermal tolerance of *A. cervicornis* corals and found evidence supporting a potential benefit of moderate ammonium enrichment. Such enhanced thermal resistance was likely mediated by poised transcriptional states established during the initial priming exposure which facilitated earlier cellular defenses upon detection of recurrent stress. Additionally, the transcriptional resilience observed in primed corals indicated a higher capacity for re-establishing homeostasis under thermal stress. These results highlight the importance of high frequency monitoring to fully characterize the dynamics of the primed transcriptional response and provide guidance on key biological processes which should be further investigated for their role in stress priming. Future work should also evaluate epigenetic mechanisms that may be responsible for maintaining primed transcriptional states without constitutive upregulation.

## Methods

### Experimental Design

Ramets of three distinct genets of *Acropora cervicornis* corals were prepared from Coral Restoration Foundation (CRF)’s nursery stock in Tavernier, Florida Keys (**Text S1**) and transported to the Keys Marine Laboratory (KML) on April 4, 2022 for the experiment. Following post-transport acclimation, corals were distributed to one of four conditions for the priming phase: ambient (control), temperature, nutrients, and combined (**Figure S1**), with full tank design and control systems described in **Text S2** and **Figure S2**. The duration of the priming and recovery phases was selected to model the naturally occurring “protective” temperature profile recorded on the Great Barrier Reef, which reduced coral bleaching relative to the “single bleaching” and “repetitive bleaching” trajectories ^19^. The magnitude of nutrient enrichment was selected within ecologically-relevant levels for the study location ^54^, with additional dosing to account for nutrient assimilation ^34^ (**Text S2**).

Within the primed treatments (temperature, nutrients, and combined), experimental conditions were gradually adjusted during the first three days of the priming phase (**Figure S3A-B**). In the elevated temperature treatments, temperature was increased (0.2 □/ hr up to 1 □/ d) from the ambient (26 □) to approximately two degrees below the bleaching threshold (29 □). In the elevated nutrient treatments, ammonium (NH4) concentrations were increased by 2 μM each day up to 6 μM above ambient. Maximum priming conditions were maintained for four additional days. On the eighth day of priming, the temperature was decreased to 26 □(0.5 □/ hr), and nutrients were returned to ambient levels through gradual water change (**Table S1**). Upon return to ambient conditions, a subset of coral fragments (4 replicates of each genotype and treatment, n=48) were flash frozen in liquid nitrogen, split into two pieces (excluding the apical tips), and preserved (stored at −80 □) for physiological and molecular analyses. Corals were then moved to 500 L tanks with constant, vigorous flow-through of ambient seawater and allowed to recover for eight days. On the eighth day, the sampling procedure was repeated on an additional 48 coral fragments (3 genotypes * 4 treatments * 4 replicates). Additionally, non-invasive measures of coral metabolism (respirometry at ambient temperature) were conducted after the priming phase (post-priming), and after five (mid-recovery) and eight (full recovery) days of recovery.

To evaluate changes in thermal tolerance induced by pre-exposure, primed corals and a subset of corals previously in the ambient treatment (now referred to as naive) were subjected to an acute thermal challenge (**Figure S1 and S3C**). Coral fragments were returned to the 150 L tanks where the water temperature was controlled using the previously-described system (**Figure S2**). During the thermal challenge, the temperature within the heated treatments was increased over 7 hr to a maximum goal temperature of 33 □. This maximum was held for 5 hr then the temperature was decreased to 26 □over 5 hr. Rather than emulating a naturally occurring marine heatwave, the acute temperature regime was selected to facilitate rapid thermal tolerance comparisons between treatment groups ^72^ and identification of molecular responses to heat at a high temporal resolution ^73^. To this end, a subset of coral fragments (3 genotypes * 5 treatments (includes addition of naive treatment) * 3 replicates) were preserved every 2-5 hours during heating for molecular analysis (total of 6 time points: challenge C1-C6). After a total of 72 hours from the onset of heating (post-challenge), a final subset of 60 coral fragments (3 genotypes * 5 treatments * 4 replicates) was preserved for physiological analyses.

### Coral Symbiosis, Metabolism, and Thermal Tolerance

At the post-priming, recovery, and post-challenge time points, coral symbiotic status was evaluated by quantifying algal symbiont density and chlorophyll concentration. One-half of each coral sample was airbrushed with sterile, ice-cold 1X PBS (phosphate-buffered saline) to separate tissue from the skeleton. Following 1 minute of homogenization with a VWR® 200 Homogenizer on ice, two 1 ml aliquots of the tissue slurry were prepared. The aliquots were centrifuged at 13,000 g for 3 min at 4 □to separate coral host and symbiont cells. The symbiont pellet of one aliquot was resuspended with 1 ml 0.01% SDS, and 900 μl of the symbiont solution was passed through a 40 μm FloMi cell strainer. The filtered symbiont solution was then fixed by adding 100 μl 10% phosphate buffered formalin. Three technical replicate aliquots were diluted (1:5) with 0.01% SDS and sent to Old Dominion University’s Center for Bioelectrics Flow Cytometry Core for flow cytometry analysis. Cell counts were standardized to total slurry volume and surface area (cells/cm^2^). The surface area of the skeleton was quantified via single wax dipping ^74^.

Chlorophyll was extracted from the symbiont pellet in the second aliquot with 1 ml of 100% acetone over a 24 hr incubation at 4 □in the dark. Chlorophyll extraction tubes were vortexed at maximum speed for 30 sec prior to and after the incubation. Extraction tubes were centrifuged at 13,000 g for 3 min at 4 □to pellet cellular debris, and 175 μl of the supernatant was pipetted in triplicate on UV clear 96-well plates (Greiner UV-Star®). Absorbance was measured at 630 nm, 663 nm, and 750 nm on an Accuris SmartReader 96 plate reader. Chlorophyll-a and c2 concentrations were calculated based on equations developed for dinoflagellates ^75^, accounting for path length ^76^, and standardized to total slurry volume and surface area (μg/cm^2^).

At the post-priming, mid-recovery, and recovery time points, the status of the coral holobiont metabolism was quantified via respirometry. Rates of dark respiration (R), net photosynthesis (Pnet), and gross photosynthesis (Pgross) were calculated from changes in dissolved O_2_ concentration as in ^77^. Recorded [O_2_] values were corrected for temperature and salinity (Witrox system). Slopes of deltaO_2_/t (ng O_2_/L/s) in the dark (R) and in the light (Pnet) were calculated from linear sections of each trace by least squares regression after at least 10 minutes under each light setting. During the light incubation, the chamber tank was illuminated with a photosynthesis-saturating irradiance of 350-450 μmol photons/m^2^/s via white LED lamps. Metabolic rates were corrected for the combined effects of instrument drift and microbial metabolism from the seawater, scaled to chamber volume (L), and normalized to coral tissue surface area (ng O_2_/s/cm^2^). Gross photosynthesis (Pgross) was calculated as Pnet + |R| and the ratio of gross photosynthesis to respiration was calculated (Pg:R).

Additionally, 48 hours into the thermal challenge, corals in the heated treatments began to show signs of tissue sloughing, which may be a common response of this species under thermal stress ^41^. Therefore, the percentage of surface area displaying complete loss of pigmentation and/or tissue loss (the combination of either case hereafter referred to as “tissue loss”) was estimated for each coral fragment every 1-3 hrs between hours 48-72 of the thermal challenge to characterize the holobiont response to this acute heat exposure.

### DNA and RNA Extraction and Gene Expression

The second half of each preserved coral sample was used for DNA/RNA co-extraction. Samples were ground into a homogeneous powder with a mortar, pestle, and liquid nitrogen, then 300 mg of powder was further homogenized with bead beating (pre-filled Zymo DNA/RNA Shield and 2.0 mm BashingBeads tubes) for two 1 min rounds (Biospec Products), alternated with 1 min on ice. Homogenized samples were further digested with Proteinase K (45 μl Proteinase K and 90 μl Proteinase K Digestion Buffer (Zymo) for 30 min at room temperature) and DNA/RNA Digestion Buffer (Zymo) with two rounds of centrifugation to remove cellular debris prior to extraction (6,700 g for 2 min at 4 □ and 12,000 g for 2 min at 4 □, prior to and following Proteinase K digestion, respectively). DNA and RNA were subsequently extracted with the Zymo Quick-DNA/RNA Miniprep Plus Kit following the manufacturer’s instructions, with the addition of one vacufuge step on an Eppendorf Vacufuge Plus at V-AL for 2 min prior to elution to remove volatile contaminants. DNA and RNA quantity and quality were confirmed with a NanoVue spectrophotometer, Qubit dsDNA and RNA broad range assay kits (Invitrogen™), and gel electrophoresis. Additionally, RNA integrity was confirmed for a subset of samples with a BioAnalyzer Agilent RNA 6000 Pico chip at the Florida International University DNA Core Facility.

DNA from all priming treatments at the post-priming and recovery time points was used to characterize the algal symbiont community. Extracted DNA was standardized to 3.5 ng/μl and amplified in triplicate PCR reactions, which included 1 μl of template DNA, 0.5 μl each of SYM_Var forward and reverse primers ^78^, 12.5 μl of Phusion™ High-Fidelity PCR Master Mix (Thermo Scientific™), and 10.5 μl molecular-grade water. PCR reaction steps followed an initial denaturing step at 95 □for 3 min, 35 cycles of denaturing (95 □for 30 s), annealing (52 □for 30 s), and extension (72 □for 30 s), and a final extension step at 72 □for 2 min. PCR products were pooled across triplicate reactions, and 20 μl was submitted for ITS2 library preparation and sequencing (paired-end, 250 bp read length, MiSeq) at the University of Rhode Island INBRE Molecular Informatics Core. Raw sequence reads were processed through the SymPortal platform to identify ITS2 sequence variants and profiles ^79^.

RNA from all treatments at the post-priming, recovery, and final challenge (C6) time points was sequenced to compare transcriptomic responses across priming stressors. Additionally, RNA from the temperature-primed, naive, and ambient treatments at all challenge time points (C1-C6) was sequenced to characterize the temporal dynamics of the primed response to heat stress. Extracted RNA was standardized to 40 ng/μl, and 3’ TagSeq library preparation and sequencing (standard coverage, NovaSeq 6000 SR100) was performed by the Genomic Sequencing and Analysis Facility at UT Austin, Center for Biomedical Research Support (RRID# SCR_021713). Quality assessment of the raw and trimmed reads was performed with FastQC. Raw reads were quality trimmed with BBDUK ^80^ using standard parameters, and resulting trimmed reads were mapped to the *Acropora cervicornis* genome ^81^, with STAR ^82^ and quantified into gene count matrices. To improve the mapping of 3’ Tag-Seq reads, the reference GFF file was modified to extend the untranslated regions of each gene 1,000 bp in each direction. Specific pipeline details can be found in the accompanying code (see Data Accessibility Statement).

### Data Analysis

#### Experimental Conditions

Water temperature (□) for each tank throughout the experiment was averaged over 10-minute intervals. Generalized additive models (GAM) (mgcv::gam, v1.9-0) ^83^ were used to model elevated water temperature (□) and ammonium concentration (μM) over time in reference to the ambient (control) treatment. Water temperature during the thermal challenge was compared between heated and control treatments using the same approach. Degree heating days were calculated for the priming and challenge phases using the cumulative temperature differential between the heated and ambient water temperatures. Similarly, cumulative nutrient enrichment was calculated during the priming phase as the accumulated concentration differential above zero.

#### Coral Symbiosis and Metabolism

Measurements of chlorophyll-a and c2 were summed to calculate total chlorophyll (μg/cm^2^), and triplicate technical replicates of chlorophyll and symbiont density (cells/cm^2^) were averaged per sample. To assess the effect of the priming treatment on coral symbiosis, linear mixed effect models (lme4::lmer, v1.1-34) ^84^ were used for each response at the post-priming and recovery time points. Within each model, the ambient treatment was the reference, and coral genotype was accounted for as a random effect. The residuals of these and all following mixed effects models were checked with the DHARMa package (v0.4.6) ^85^. The overall effect of the priming treatment was evaluated with Type III ANOVA (analysis of variance) and quantified as partial eta-squared (effectsize::eta_squared, v0.8.6) ^86,87^. The same analysis pipeline was applied to gross photosynthesis (P_Gross_, ng O_2_/cm^2^/s), respiration (R, ng O_2_/cm^2^/s), and P_Gross_:R measured at the post-priming, mid-recovery, and recovery time points.

#### Thermal Tolerance

Thermotolerance was quantified as the retention of symbionts and chlorophyll at the end of the thermal challenge, as well as live tissue retention throughout this phase. For symbiont and chlorophyll retention, the average value for each genet in the ambient treatment was considered 100%, and the proportion retained was then calculated for each heated sample. To compare the thermal tolerance of primed versus naive corals, generalized linear mixed effects models (lme4::glmer, v1.1-34; binomial distribution) of symbiont and chlorophyll retention were used. The naive treatment was the reference, and the coral genotype was included as a random effect. The odds ratio was calculated as the exponential of the coefficient estimate for each primed treatment compared to the naive treatment.

To quantify thermal tolerance in terms of live tissue retention throughout the thermal challenge, two modeling approaches were utilized. First, the declines in live tissue over time were compared between the naive and primed treatments with generalized linear mixed effects models (binomial distribution; lme4::glmer, v1.1-34). The proportion of live tissue was modeled as a function of time plus the interaction between time and priming treatment, with a random effect of genotype. The slopes of each priming treatment were compared to the naive treatment (glmm model fixed effects) as well as each other through pairwise comparisons with Tukey’s Honest Significant Difference (HSD) corrections (emmeans::lstrends, v1.8.9) ^88,89^. Survival analysis was also performed to compare the onset of tissue loss between treatments. Cox proportional hazards models, clustered by coral genotype, were used to model the occurrence of two different thresholds of tissue loss (survival::coxph, v3.5-5) ^90^, including the onset of any visible tissue loss (> 0%) and tissue loss greater than 20% surface area as an established proxy for predicted mortality of this species ^91^. The first model compared the risk of tissue loss in all heated treatments (*i.e.,* primed and naive treatments) to that of the ambient control treatment, while the second only included heated treatments to compare the likelihood of survival between the primed and the naive treatments. Pairwise comparisons were also performed as previously described.

#### Molecular Analysis

The symbiont community composition was described by the relative abundance of symbiont types identified at the level of sequence variants and predicted ITS2 profiles ^79^. Permutational analysis of variance (PERMANOVA) and dispersion (PERMDISP) (vegan::adonis2 and betadisper, v2.6-4) were used to compare symbiont communities (Bray-Curtis distance) across coral genotypes, priming treatments, and time points (post-priming and recovery) ^92^. The effect size of each predictor was quantified as partial omega squared (MicEco::adonis_OmegaSq, v0.9.19). Post-hoc pairwise comparisons of variance (pairwiseAdonis::pairwise.adonis, v0.4.1) and dispersion (TukeyHSD) were conducted for significant predictors.

To characterize transcriptomic responses to stress priming, differential expression analysis was conducted with the DESeq2 package (v1.40.2) ^93^. Metadata fields were derived to encode experimental structure in the gene count matrix, including priming conditions, phase (Priming [PP], Recovery [RE], Challenge [C1–C6]), and treatment status. Genes with low expression were filtered within each analysis subset by retaining only features with >10 counts in at least three samples (minimum replicate number). For differential expression models, an initial assessment indicated minimal plate-associated variation, and the plate was therefore omitted from downstream models, with genotype included as a covariate. Differential expression was tested with negative binomial generalized linear models and Benjamini–Hochberg FDR control ^94,95^. Priming effects were evaluated by subsetting to PP and RE time points and fitting *∼ Genotype + time point * Treatment* with PP and Ambient as reference levels, enabling treatment contrasts at each time point and time point contrasts within each treatment. End-of-challenge effects were assessed at C6 using *∼ Genotype + Treatment*. To quantify priming effects across the thermal challenge (C1–C6), challenge-phase samples were analyzed with *∼ Genotype + time point * Treatment* (C1 and Ambient as references), and time point-specific contrasts were constructed for Naive vs Control, Primed vs Control, and Primed vs Naive (difference-of-differences). Log2 fold changes were shrunk for reporting using *apeglm* for single-coefficient contrasts and *ashr* for linear (multi-coefficient) contrasts ^96^, while unshrunken statistics were retained for inference. Functional interpretation used a custom gene-to-GO mapping and clusterProfiler::enricher (v4.8.3; ^97^ with the appropriate tested-gene universe, restricting to Biological Process terms and controlling FDR (q < 0.05).

To identify coordinated transcriptional programs during the challenge, variance-stabilized expression (DESeq2 VST; blind = FALSE) from challenge-phase samples was used for weighted gene coexpression network analysis (WGCNA; ^98^) after retaining the most variable genes (top ∼70% by variance). A signed network was constructed using biweight midcorrelation, soft-threshold power selection, and module detection with blockwiseModules (minimum module size 30; merge cut height 0.25). Module eigengenes were correlated with encoded traits (challenged status, primed/naive indicators, and time), and eigengene trajectories were summarized as mean ± SE across time points within treatment groups. Module gene sets were functionally annotated by GO enrichment as described above.

## Acknowledgments

We are thankful to the Coral Restoration Foundation for sharing the *Acropora* genets used in the present study and for logistic support. This work would not have been possible without the dedicated help from members of FIU’s Environmental Epigenetics Lab and Marine Sciences Program. This is contribution #TBD from the Institute of Environment at Florida International University.

## Author contributions

S.H.: conceived study, acquired funding, performed fieldwork, developed experiments, analyzed data, wrote manuscript. H.M.: conceived study, acquired funding, performed fieldwork, analyzed data, edited manuscript. J.R.C.: analyzed data, edited manuscript. J.E.L.: conceived study, acquired funding, supervised study, wrote manuscript.

## Data availability

All raw data files and associated analysis scripts and code are available on the Environmental Epigenetics Lab GitHub at https://github.com/eelabfiu/CSHORE_priming

