## Supplemental materials for "Combined-stress priming alters physiology, transcriptional dynamics, and acute thermal resistance in *Acropora cervicornis* corals"

**Supplemental Information**

**Supplemental Text**

**Text S1.** Coral Species and Ramet Preparation

Ramets of three distinct genets of *Acropora cervicornis* corals (AC032, AC062, and AC112) were prepared from Coral Restoration Foundation (CRF)’s offshore nursery stock in Tavernier, Florida Keys. Ramets, 8-10 cm in length, were prepared, secured to nylon bolts with marine epoxy (AquaMend), then randomly distributed and affixed across egg crate grids. Grids were attached to the CRF nursery tree structures for ramets to heal for one month.

**Text S2.** Experimental Design and Aquarium Set Up

The experiment was conducted in the Keys Marine Laboratory (KML) outdoor, shaded seawater tank system. Powerheads (Aqueon, 3.6 kL/hr) were added to each tank for water circulation. Water flow and circulation were paused for one hour every other day when corals were fed Reef-Roids (Polylab) following the manufacturer's instructions. The salinity of each tank was measured with a refractometer each day. The temperature of each tank was monitored with HOBO pendant loggers (Onset MX2201) every minute. On April 4, 2022, corals were transported to KML and acclimated for five days in four 500 L tanks with constant flow-through of ambient seawater plus three powerheads. Corals were then distributed to one of four conditions for the priming phase: ambient (control), temperature, nutrients, and combined (**Figure S1**) across twelve 150 L tanks. During the priming phase, the water temperature of each tank was controlled independently using the Neptune APEX system (**Figure S2**). Elevated nutrient concentrations were established in the nutrients and combined treatments through the addition of a 0.3M NH_4_Cl stock solution to reach the goal concentration of ammonium. Additionally, two APEX DOS modules added 0.2 mL of 6.25 mM NH_4_Cl every 12 min to account for nutrient assimilation. Throughout the priming phase, seawater flow-through was terminated and replaced with daily water changes. Corals were shuffled within tanks of the same treatment to maintain experimental conditions during tank maintenance. Water samples were collected from six tanks each day during the priming phase and filtered through a 0.22 μm nylon filter (CELLTREAT) before freezing. Ammonium concentration was measured colorimetrically at the Florida International University CAChE Nutrient Analysis Core Facility (NELAC Certified Analysis, Certificate# E76930-16) following EPA method 349.0 on a four-channel Rapid Flow Analyzer (Astoria-Pacific).

**Supplemental Tables**

**Table S1. Experimental conditions.** Parameters describing exposure to elevated ammonium and temperature in the priming and thermal challenge phases (**Figure S1 and S2**). Averages are across tanks (and days for ambient and maximum value statistics) ± standard error.

|  | **Ammonium** | **Temperature** | |
| --- | --- | --- | --- |
|  | *Priming Phase* | *Priming Phase* | *Thermal Challenge* |
| **Ambient** | 0.520 ± 0.157 (μM) | 25.999 ± 0.003 (℃) | 26.093 ± 0.007 (℃) |
| **Maximum Value** | 3.815 ± 0.204 (μM) | 28.850 ± 0.008 (℃) | 32.891 ± 0.020 (℃) |
| **Ramp Rate Increase** | 3.745 ± 0.193 (μM/day) | 0.993 ± 0.016 (℃/day) | 0.946 ± 0.019 (℃/hr) |
| **Duration of Maximum** | 4 days | 4 days | 5 hours |
| **Ramp Rate Decrease** | 1.265 ± 0.140 (μM/day) | 2.952 ± 0.069 (℃/day) | 1.383 ± 0.024 (℃/hr) |
| **Cumulative Exposure** | 16.436 ± 1.292 (μM * day) | 17.222 ± 0.277 (℃ * day) | 3.251 ± 0.345 (℃ * day) |

**Table S2. Model results comparing experimental treatments.** Results of generalized additive models (GAM) of **A)** temperature (℃) and **B)** ammonium concentration (μM) throughout the priming phase (**Figure S3A-B**) and **C)** temperature (℃) during the thermal challenge (**Figure S3C**). For each parameter, each treatment is smoothed over time and the estimated degrees of freedom (DF), F value, and p value (bold: p < 0.05) are listed for each smooth term. The adjusted R^2^ and percent of deviance explained by the full model are shown.

| **A) Temperature Priming** | | | |
| --- | --- | --- | --- |
| **Smooth Terms** | **Estimated DF** | **F** | **p** |
| Ambient | 8.156 | 16.973 | **< 0.001** |
| Temperature | 8.971 | 1,874.91 | **< 0.001** |
| Nutrients | 1 | 1.449 | 0.229 |
| Combined | 8.983 | 1,566.71 | **< 0.001** |
| **Adj. R-sq =** | 0.946 | **Deviance explained =** | 94.60% |
| **B) Nutrient Priming** | | | |
| **Smooth Terms** | **Estimated DF** | **F** | **p** |
| Ambient | 2.561 | 4.119 | **0.027** |
| Nutrients | 2.43 | 6.499 | **0.005** |
| Combined | 2.685 | 12.497 | **< 0.001** |
| **Adj. R-sq =** | 0.83 | **Deviance explained =** | 86.60% |
| **C) Thermal Challenge** | | | |
| **Smooth Terms** | **Estimated DF** | **F** | **p** |
| Control | 8.336 | 10.8 | **< 0.001** |
| Heated | 8.981 | 2,374.30 | **< 0.001** |
| **Adj. R-sq =** | 0.942 | **Deviance explained =** | 94.20% |

**Table S3. Model results comparing bleaching metrics across treatments.** Results of linear mixed effects models of chlorophyll concentration (μg/cm^2^) at the **A)** post-priming and **B)** recovery time points and **C)** post-priming symbiont density (cells/cm^2^) (**Figure S4**). The coefficient estimate of each primed treatment, in comparison to the ambient treatment, along with the standard error (SE), degrees of freedom (DF), t value, and p value (bold: p < 0.05) are listed. Additionally, the overall treatment effect (Analysis of Variance, ANOVA) is shown for each metric and time point, including sum of squares (SS), mean squares (MS), numerator and denominator DF, F value, p value, and effect size (partial eta-squared).

| **A) Chlorophyll Concentration Post-Priming** | | | | | |
| --- | --- | --- | --- | --- | --- |
| **Predictor** | **Estimate** | **SE** | **DF** | **t** | **p** |
| Intercept | 4.184 | 0.369 | 3.447 | 11.328 | **0.001** |
| Temperature | -0.933 | 0.287 | 40.014 | -3.247 | **0.002** |
| Nutrients | -0.458 | 0.287 | 40.014 | -1.594 | 0.119 |
| Combined | -0.637 | 0.294 | 40.046 | -2.164 | **0.037** |
| **ANOVA** | **SS / MS** | **NumDF / DenDF** | **F** | **p** | **Partial Eta2** |
| Treatment | 5.206 / 1.735 | 3 / 40.016 | 3.666 | **0.02** | 0.216 |
| **B) Chlorophyll Concentration End of Recovery** | | | | | |
| **Predictor** | **Estimate** | **SE** | **DF** | **t** | **p** |
| Intercept | 4.505 | 0.262 | 5.944 | 17.19 | **< 0.001** |
| Temperature | -0.905 | 0.28 | 42 | -3.236 | **0.002** |
| Nutrients | -0.852 | 0.28 | 42 | -3.046 | **0.004** |
| Combined | -0.676 | 0.28 | 42 | -2.415 | **0.02** |
| **ANOVA** | **SS / MS** | **NumDF / DenDF** | **F** | **p** | **Partial Eta2** |
| Treatment | 6.269 / 2.09 | 3 / 42 | 4.448 | **0.008** | 0.241 |
| **C) Symbiont Density Post-Priming** | | | | | |
| **Predictor** | **Estimate** | **SE** | **DF** | **t** | **p** |
| Intercept | 1.603 | 0.141 | 9.505 | 11.336 | **< 0.001** |
| Temperature | -0.048 | 0.167 | 40.093 | -0.286 | 0.776 |
| Nutrients | -0.041 | 0.167 | 40.093 | -0.242 | 0.81 |
| Combined | 0.151 | 0.171 | 40.214 | 0.884 | 0.382 |
| **ANOVA** | **SS / MS** | **NumDF / DenDF** | **F** | **p** | **Partial Eta2** |
| Treatment | 0.291 / 0.097 | 3 / 40.1 | 0.604 | 0.616 | 0.043 |

**Table S4. Model results comparing metabolic metrics across treatments.** Results of linear mixed effects models of the ratio of gross photosynthesis to respiration (P_Gross_ : R), gross photosynthesis (ng O_2_/cm^2^/s), and respiration (ng O_2_/cm^2^/s) at the **A)** post-priming and **B)** mid-recovery, and **C)** end of recovery time points (**Figure S4**). The coefficient estimate of each primed treatment, in comparison to the ambient treatment, along with the standard error (SE), degrees of freedom (DF), t value, and p value (bold: p < 0.05) are listed. Additionally, the overall treatment effect (Analysis of Variance, ANOVA) is shown for each metric and time point, including sum of squares (SS), mean squares (MS), numerator and denominator DF, F value, p value, and effect size (partial eta-squared).

| **A) Post-Priming** | | | | | | |
| --- | --- | --- | --- | --- | --- | --- |
| **Response** | **Predictor** | **Estimate** | **SE** | **DF** | **t** | **p** |
| P:R | Intercept | 4.012 | 0.323 | 11.092 | 12.418 | **< 0.001** |
|  | Temperature | 1.087 | 0.53 | 63.613 | 2.049 | **0.045** |
|  | Nutrients | 0.962 | 0.475 | 63.329 | 2.023 | **0.047** |
|  | Combined | 1.542 | 0.485 | 64.656 | 3.178 | **0.002** |
| Photosynthesis | Intercept | 8.82 | 0.96 | 3.021 | 9.185 | **0.003** |
|  | Temperature | 0.955 | 0.874 | 63.122 | 1.093 | 0.279 |
|  | Nutrients | 0.332 | 0.782 | 63.066 | 0.424 | 0.673 |
|  | Combined | 0.811 | 0.804 | 63.332 | 1.01 | 0.317 |
| Respiration | Intercept | 2.343 | 0.175 | 4.314 | 13.411 | **< 0.001** |
|  | Temperature | -0.364 | 0.209 | 63.242 | -1.743 | 0.086 |
|  | Nutrients | -0.432 | 0.187 | 63.132 | -2.306 | **0.024** |
|  | Combined | -0.559 | 0.192 | 63.659 | -2.909 | **0.005** |
| **Response** | **ANOVA** | **SS / MS** | **NumDF / DenDF** | **F** | **p** | **Partial Eta2** |
| P:R | Treatment | 25.515 / 8.505 | 3 / 64.128 | 3.783 | **0.015** | 0.15 |
| Photosynthesis | Treatment | 10.183 / 3.394 | 3 / 63.221 | 0.558 | 0.645 | 0.026 |
| Respiration | Treatment | 3.549 / 1.183 | 3 / 63.438 | 3.398 | **0.023** | 0.138 |
| **B) Mid-Recovery** | | | | | | |
| **Response** | **Predictor** | **Estimate** | **SE** | **DF** | **t** | **p** |
| P:R | Intercept | 5.938 | 0.486 | 80 | 12.215 | **< 0.001** |
|  | Temperature | 0.323 | 0.842 | 80 | 0.383 | 0.703 |
|  | Nutrients | 1.613 | 0.862 | 80 | 1.871 | 0.065 |
|  | Combined | -0.396 | 0.721 | 80 | -0.549 | 0.585 |
| Photosynthesis | Intercept | 9.267 | 0.81 | 80 | 11.442 | **< 0.001** |
|  | Temperature | 0.668 | 1.403 | 80 | 0.476 | 0.635 |
|  | Nutrients | -0.564 | 1.436 | 80 | -0.393 | 0.695 |
|  | Combined | 1.073 | 1.201 | 80 | 0.893 | 0.375 |
| Respiration | Intercept | 1.686 | 0.169 | 3.552 | 9.973 | **0.001** |
|  | Temperature | -0.038 | 0.184 | 78.434 | -0.206 | 0.837 |
|  | Nutrients | -0.401 | 0.187 | 78.081 | -2.14 | **0.036** |
|  | Combined | 0.263 | 0.158 | 78.437 | 1.668 | 0.099 |
| **Response** | **ANOVA** | **SS / MS** | **NumDF / DenDF** | **F** | **p** | **Partial Eta2** |
| P:R | Treatment | 38.221 / 12.74 | 3 / 80 | 1.797 | 0.154 | 0.063 |
| Photosynthesis | Treatment | 29.95 / 9.983 | 3 / 80 | 0.507 | 0.678 | 0.019 |
| Respiration | Treatment | 3.93 / 1.31 | 3 / 78.426 | 3.917 | **0.012** | 0.13 |
| **C) End of Recovery** | | | | | | |
| **Response** | **Predictor** | **Estimate** | **SE** | **DF** | **t** | **p** |
| P:R | Intercept | 5.861 | 0.576 | 10.872 | 10.173 | **< 0.001** |
|  | Temperature | -0.915 | 0.819 | 24.712 | -1.118 | 0.274 |
|  | Nutrients | 0.811 | 0.85 | 25.043 | 0.954 | 0.349 |
|  | Combined | 0.384 | 0.888 | 24.469 | 0.432 | 0.669 |
| Photosynthesis | Intercept | 8.316 | 1.474 | 3.756 | 5.641 | **0.006** |
|  | Temperature | 0.377 | 1.305 | 24.227 | 0.289 | 0.775 |
|  | Nutrients | 0.562 | 1.36 | 24.318 | 0.414 | 0.683 |
|  | Combined | 2.421 | 1.409 | 24.119 | 1.718 | 0.099 |
| Respiration | Intercept | 1.531 | 0.289 | 4.994 | 5.3 | **0.003** |
|  | Temperature | 0.277 | 0.317 | 24.204 | 0.873 | 0.391 |
|  | Nutrients | -0.038 | 0.33 | 24.377 | -0.114 | 0.91 |
|  | Combined | 0.196 | 0.343 | 24.012 | 0.57 | 0.574 |
| **Response** | **ANOVA** | **SS / MS** | **NumDF / DenDF** | **F** | **p** | **Partial Eta2** |
| P:R | Treatment | 12.167 / 4.056 | 3 / 24.673 | 1.431 | 0.258 | 0.148 |
| Photosynthesis | Treatment | 23.008 / 7.669 | 3 / 24.191 | 1.079 | 0.377 | 0.118 |
| Respiration | Treatment | 0.526 / 0.175 | 3 / 24.144 | 0.416 | 0.743 | 0.049 |

**Table S5. Model results comparing symbiont communities across treatments.** Results of permutational analyses of variance (PERMANOVA) and dispersion (PERMDISP) comparing symbiont community composition at the level of **A)** sequence variants and **B)** ITS2 profiles across genotypes, treatments, and timepoints. For each predictor, degrees of freedom (DF), sum of squares (SS), F value, effect size (partial omega-squared), and p values are shown, along with PERMDISP p values when appropriate (bold: p < 0.05).

| **A) Sequence Variants** | | | | | | |
| --- | --- | --- | --- | --- | --- | --- |
| **Predictor** | **DF** | **SS** | **F** | **Partial Omega2** | **p** | **p Dispersion** |
| Genotype | 2 | 0.01 | 40.691 | 0.524 | **0.001** | **0.015** |
| Treatment | 3 | 0 | 1.309 | 0.013 | 0.25 | *NA* |
| Timepoint | 1 | 0 | 4.204 | 0.043 | **0.028** | 0.697 |
| Treatment:Timepoint | 3 | 0 | 0.405 | -0.025 | 0.854 | *NA* |
| Genotype:Treatment | 6 | 0 | 0.686 | -0.027 | 0.71 | *NA* |
|  | **DF** | **SS** |  | **DF** | **SS** |  |
| **Residual** | 56 | 0.007 | **Total** | 71 | 0.018 |  |
| **Pairwise Contrast** | **DF** | **SS** | **R2** | **F** | **p** | **p Dispersion** |
| AC032 vs AC062 | 1 | 0.007 | 0.546 | 55.211 | **0.001** | **0.011** |
| AC032 vs AC112 | 1 | 0.006 | 0.65 | 85.28 | **0.001** | 0.308 |
| AC062 vs AC112 | 1 | 0.001 | 0.159 | 8.679 | **0.001** | 0.297 |
| **B) ITS2 Profiles** | | | | | | |
| **Predictor** | **DF** | **SS** | **F** | **Partial Omega2** | **p** | **p Dispersion** |
| Genotype | 2 | 0.083 | 0.955 | -0.001 | 0.652 | *NA* |
| Treatment | 3 | 0.153 | 1.167 | 0.007 | 0.471 | *NA* |
| Timepoint | 1 | 0.014 | 0.318 | -0.01 | 0.942 | *NA* |
| Treatment:Timepoint | 3 | 0.042 | 0.318 | -0.029 | 0.975 | *NA* |
| Genotype:Treatment | 6 | 0.139 | 0.53 | -0.041 | 0.921 | *NA* |
|  | **DF** | **SS** |  | **DF** | **SS** |  |
| **Residual** | 56 | 2.445 | **Total** | 71 | 2.875 |  |

**Table S6. Results of Gene Ontology (GO) functional enrichment analysis after priming.** Result of overrepresentation analysis of gene ontology (GO) terms of biological processes for genes **A)** upregulated and **B)** downregulated in both temperature- and combined-primed corals at the post-priming time point (**Figure 1**). For each GO term, the gene ratio (significant genes / total genes in term), term ratio (total genes in term / total genes), p value (bold: p < 0.05), and q value are listed.

| **A) Shared Upregulated** | | | | | |
| --- | --- | --- | --- | --- | --- |
| **ID** | **Description** | **Gene Ratio** | **Term Ratio** | **p** | **q** |
| GO:0003341 | cilium movement | 7/559 | 23/12045 | **0.021** | 0.021 |
| GO:0060294 | cilium movement involved in cell motility | 5/559 | 12/12045 | **0.021** | 0.021 |
| GO:0007266 | Rho protein signal transduction | 7/559 | 27/12045 | **0.021** | 0.021 |
| GO:0060271 | cilium assembly | 12/559 | 78/12045 | **0.021** | 0.021 |
| **B) Shared Downregulated** | | | | | |
| **ID** | **Description** | **Gene Ratio** | **Term Ratio** | **p** | **q** |
| GO:0032970 | regulation of actin filament-based process | 9/699 | 20/12045 | **< 0.001** | 0 |
| GO:0060415 | muscle tissue morphogenesis | 6/699 | 10/12045 | **0.001** | 0.001 |
| GO:1903566 | positive regulation of protein localization to cilium | 6/699 | 12/12045 | **0.004** | 0.003 |
| GO:0010638 | positive regulation of organelle organization | 6/699 | 14/12045 | **0.007** | 0.006 |
| GO:0006749 | glutathione metabolic process | 7/699 | 20/12045 | **0.007** | 0.006 |
| GO:0051240 | positive regulation of multicellular organismal process | 9/699 | 35/12045 | **0.009** | 0.008 |
| GO:0051493 | regulation of cytoskeleton organization | 7/699 | 22/12045 | **0.01** | 0.009 |
| GO:0060341 | regulation of cellular localization | 8/699 | 32/12045 | **0.02** | 0.017 |
| GO:0051234 | establishment of localization | 24/699 | 197/12045 | **0.02** | 0.018 |
| GO:0032880 | regulation of protein localization | 9/699 | 42/12045 | **0.02** | 0.018 |
| GO:0032386 | regulation of intracellular transport | 5/699 | 13/12045 | **0.02** | 0.018 |
| GO:0010564 | regulation of cell cycle process | 8/699 | 34/12045 | **0.02** | 0.018 |
| GO:0042127 | regulation of cell population proliferation | 10/699 | 54/12045 | **0.031** | 0.027 |
| GO:0051098 | regulation of binding | 6/699 | 22/12045 | **0.038** | 0.033 |
| GO:0032501 | multicellular organismal process | 24/699 | 213/12045 | **0.039** | 0.034 |
| GO:2000026 | regulation of multicellular organismal development | 9/699 | 48/12045 | **0.041** | 0.035 |
| GO:0043393 | regulation of protein binding | 4/699 | 10/12045 | **0.042** | 0.036 |
| GO:0043666 | regulation of phosphoprotein phosphatase activity | 4/699 | 10/12045 | **0.042** | 0.036 |
| GO:0042981 | regulation of apoptotic process | 15/699 | 113/12045 | **0.044** | 0.039 |
| GO:0010941 | regulation of cell death | 5/699 | 17/12045 | **0.044** | 0.039 |
| GO:0032879 | regulation of localization | 14/699 | 103/12045 | **0.044** | 0.039 |
| GO:0034976 | response to endoplasmic reticulum stress | 4/699 | 11/12045 | **0.044** | 0.039 |
| GO:0048609 | multicellular organismal reproductive process | 4/699 | 11/12045 | **0.044** | 0.039 |
| GO:0051014 | actin filament severing | 4/699 | 11/12045 | **0.044** | 0.039 |
| GO:0051239 | regulation of multicellular organismal process | 15/699 | 116/12045 | **0.044** | 0.039 |
| GO:0031122 | cytoplasmic microtubule organization | 7/699 | 34/12045 | **0.044** | 0.039 |
| GO:0080134 | regulation of response to stress | 8/699 | 43/12045 | **0.044** | 0.039 |
| GO:0006418 | tRNA aminoacylation for protein translation | 5/699 | 18/12045 | **0.044** | 0.039 |
| GO:0030198 | extracellular matrix organization | 8/699 | 44/12045 | **0.049** | 0.043 |

**Table S7. Model results comparing thermal tolerance across treatments.** Results of generalized linear mixed effects models of **A)** symbiont and **B)** chlorophyll retention after acute heating as well as **C-D)** live tissue retained over time and cox proportional hazards models of the risks of **E)** tissue loss and **F)** predicted mortality throughout the thermal challenge (**Figures 2 and S6**). In addition to coefficient estimates, standard error (SE), z value, and p value (bold: p < 0.05), odds and hazards ratios are shown for panels **A-B** and **E-F** as the exponential of the coefficient. Post-hoc pairwise comparisons are shown for panels **C-D**.

| **A) Symbiont Retention** | | | | | | |
| --- | --- | --- | --- | --- | --- | --- |
| **Predictor** | **Reference** | **Estimate** | **Odds Ratio** | **SE** | **z** | **p** |
| Intercept | Naive | -0.767 | 0.464 | 0.696 | -1.103 | 0.27 |
| Temperature | Naive | 1.468 | 4.342 | 0.871 | 1.686 | 0.092 |
| Nutrients | Naive | 1.08 | 2.944 | 0.816 | 1.323 | 0.186 |
| Combined | Naive | 1.882 | 6.565 | 0.856 | 2.199 | **0.028** |
| **B) Chlorophyll Retention** | | | | | | |
| **Predictor** | **Reference** | **Estimate** | **Odds Ratio** | **SE** | **z** | **p** |
| Intercept | Naive | -1.115 | 0.328 | 0.734 | -1.519 | 0.129 |
| Temperature | Naive | 1.426 | 4.161 | 0.877 | 1.625 | 0.104 |
| Nutrients | Naive | 1.435 | 4.2 | 0.848 | 1.693 | 0.091 |
| Combined | Naive | 1.889 | 6.613 | 0.858 | 2.202 | **0.028** |
| **C) Live Tissue All Treatments** | | | | | | |
| **Predictor** | **Reference** | **Estimate** | **SE** | **z** | **p** |  |
| Intercept | Ambient | 4.8 | 0.309 | 15.526 | **< 0.001** |  |
| Time | Ambient | 0.271 | 0.085 | 3.201 | **0.001** |  |
| Time : Naive | Ambient | -0.35 | 0.086 | -4.078 | **< 0.001** |  |
| Time : Temperature | Ambient | -0.328 | 0.086 | -3.823 | **< 0.001** |  |
| Time : Nutrients | Ambient | -0.325 | 0.086 | -3.795 | **< 0.001** |  |
| Time : Combined | Ambient | -0.322 | 0.086 | -3.757 | **< 0.001** |  |
| **Pairwise Contrast** | | **Estimate** | **SE** | **z Ratio** | **p** |  |
| Ambient vs Naive | | 0.35 | 0.086 | 4.078 | **< 0.001** |  |
| Ambient vs Temperature | | 0.328 | 0.086 | 3.823 | **0.001** |  |
| Ambient vs Nutrients | | 0.325 | 0.086 | 3.795 | **0.001** |  |
| Ambient vs Combined | | 0.322 | 0.086 | 3.757 | **0.002** |  |
| Naive vs Temperature | | -0.022 | 0.004 | -5.093 | **< 0.001** |  |
| Naive vs Nutrients | | -0.024 | 0.004 | -5.742 | **< 0.001** |  |
| Naive vs Combined | | -0.028 | 0.004 | -6.486 | **< 0.001** |  |
| Temperature vs Nutrients | | -0.002 | 0.005 | -0.536 | 0.984 |  |
| Temperature vs Combined | | -0.006 | 0.005 | -1.259 | 0.716 |  |
| Nutrients vs Combined | | -0.003 | 0.004 | -0.738 | 0.948 |  |
| **D) Live Tissue Heated Treatments** | | | | | | |
| **Predictor** | **Reference** | **Estimate** | **SE** | **z** | **p** |  |
| Intercept | Naive | 38.315 | 6.276 | 6.105 | **< 0.001** |  |
| Time | Naive | -0.637 | 0.109 | -5.836 | **< 0.001** |  |
| Time : Temperature | Naive | 0.226 | 0.07 | 3.243 | **0.001** |  |
| Time : Nutrients | Naive | 0.202 | 0.065 | 3.125 | **0.002** |  |
| Time : Combined | Naive | 0.21 | 0.064 | 3.278 | **0.001** |  |
| **Pairwise Contrast** | | **Estimate** | **SE** | **z Ratio** | **p** |  |
| Naive vs Temperature | | -0.226 | 0.07 | -3.243 | **0.006** |  |
| Naive vs Nutrients | | -0.202 | 0.065 | -3.125 | **0.01** |  |
| Naive vs Combined | | -0.21 | 0.064 | -3.278 | **0.006** |  |
| Temperature vs Nutrients | | 0.024 | 0.043 | 0.544 | 0.948 |  |
| Temperature vs Combined | | 0.015 | 0.043 | 0.356 | 0.985 |  |
| Nutrients vs Combined | | -0.008 | 0.034 | -0.244 | 0.995 |  |
| **E) Tissue Loss (Tissue Loss > 0%)** | | | | | | |
| **Predictor** | **Reference** | **Estimate** | **Hazard Ratio** | **SE** | **z** | **p** |
| Naive | Ambient | 3.121 | 22.675 | 1.042 | 3.046 | **0.002** |
| Temperature | Ambient | 1.732 | 5.65 | 1.118 | 1.226 | 0.22 |
| Nutrients | Ambient | 2.119 | 8.323 | 1.081 | 2.076 | **0.038** |
| Combined | Ambient | 1.66 | 5.261 | 1.118 | 1.276 | 0.202 |
| **F) Predicted Mortality (Tissue Loss > 20%)** | | | | | | |
| **Predictor** | **Reference** | **Estimate** | **Hazard Ratio** | **SE** | **z** | **p** |
| Temperature | Naive | -1.583 | 0.205 | 0.649 | -4.568 | **< 0.001** |
| Nutrients | Naive | -0.954 | 0.385 | 0.502 | -1.874 | 0.061 |
| Combined | Naive | -1.391 | 0.249 | 0.58 | -2.602 | **0.009** |

**Table S8. Results of Gene Ontology (GO) functional enrichment analysis after thermal challenge.** Result of overrepresentation analysis of gene ontology (GO) terms of biological processes for genes **A)** upregulated and **B)** downregulated in naive corals, **C)** upregulated and **D)** downregulated in temperature-primed corals, **E)** upregulated and **F)** downregulated in nutrient-primed corals, and **E)** upregulated and **F)** downregulated in combined-primed corals at the end of acute heating in the thermal challenge (time point C6) (**Figures 3 and S7**). For each GO term, the gene ratio (significant genes / total genes in term), term ratio (total genes in term / total genes), p value (bold: p < 0.05), and q value are listed.

| **A) Induced in Naive** | | | | | |
| --- | --- | --- | --- | --- | --- |
| **ID** | **Description** | **Gene Ratio** | **Term Ratio** | **p** | **q** |
| GO:0043161 | proteasome-mediated ubiquitin-dependent protein catabolic process | 38/2220 | 88/11632 | **< 0.001** | 0 |
| GO:0006355 | regulation of DNA-templated transcription | 101/2220 | 348/11632 | **0.001** | 0.001 |
| GO:0051603 | proteolysis involved in protein catabolic process | 23/2220 | 48/11632 | **0.001** | 0.001 |
| GO:0006511 | ubiquitin-dependent protein catabolic process | 34/2220 | 89/11632 | **0.003** | 0.002 |
| GO:0007266 | Rho protein signal transduction | 15/2220 | 27/11632 | **0.003** | 0.003 |
| GO:0006357 | regulation of transcription by RNA polymerase II | 108/2220 | 395/11632 | **0.003** | 0.003 |
| GO:0003341 | cilium movement | 13/2220 | 23/11632 | **0.005** | 0.005 |
| GO:0010498 | proteasomal protein catabolic process | 9/2220 | 13/11632 | **0.007** | 0.006 |
| GO:0006906 | vesicle fusion | 12/2220 | 21/11632 | **0.007** | 0.006 |
| GO:0030433 | ubiquitin-dependent ERAD pathway | 16/2220 | 33/11632 | **0.007** | 0.006 |
| GO:0035556 | intracellular signal transduction | 45/2220 | 143/11632 | **0.012** | 0.012 |
| GO:0006888 | endoplasmic reticulum to Golgi vesicle-mediated transport | 22/2220 | 56/11632 | **0.015** | 0.015 |
| GO:0006413 | translational initiation | 12/2220 | 24/11632 | **0.025** | 0.024 |
| GO:0016567 | protein ubiquitination | 38/2220 | 123/11632 | **0.041** | 0.04 |
| **B) Repressed in Naive** | | | | | |
| **ID** | **Description** | **Gene Ratio** | **Term Ratio** | **p** | **q** |
| GO:0006260 | DNA replication | 30/3222 | 42/11632 | **< 0.001** | 0 |
| GO:0032508 | DNA duplex unwinding | 17/3222 | 22/11632 | **< 0.001** | 0 |
| GO:0055085 | transmembrane transport | 154/3222 | 403/11632 | **< 0.001** | 0 |
| GO:0007099 | centriole replication | 10/3222 | 10/11632 | **< 0.001** | 0 |
| GO:0006281 | DNA repair | 47/3222 | 97/11632 | **0.001** | 0.001 |
| GO:0000727 | double-strand break repair via break-induced replication | 10/3222 | 11/11632 | **0.002** | 0.002 |
| GO:0030198 | extracellular matrix organization | 25/3222 | 43/11632 | **0.002** | 0.002 |
| GO:0032970 | regulation of actin filament-based process | 14/3222 | 19/11632 | **0.003** | 0.002 |
| GO:0006302 | double-strand break repair | 19/3222 | 30/11632 | **0.003** | 0.003 |
| GO:0006508 | proteolysis | 111/3222 | 291/11632 | **0.003** | 0.003 |
| GO:0006412 | translation | 54/3222 | 126/11632 | **0.008** | 0.008 |
| GO:0060341 | regulation of cellular localization | 18/3222 | 31/11632 | **0.017** | 0.015 |
| GO:1902476 | chloride transmembrane transport | 17/3222 | 29/11632 | **0.019** | 0.017 |
| GO:0000723 | telomere maintenance | 13/3222 | 20/11632 | **0.021** | 0.019 |
| GO:0060415 | muscle tissue morphogenesis | 8/3222 | 10/11632 | **0.03** | 0.027 |
| GO:1903566 | positive regulation of protein localization to cilium | 9/3222 | 12/11632 | **0.03** | 0.027 |
| GO:0005975 | carbohydrate metabolic process | 53/3222 | 131/11632 | **0.032** | 0.029 |
| GO:0000724 | double-strand break repair via homologous recombination | 23/3222 | 46/11632 | **0.032** | 0.029 |
| GO:0006270 | DNA replication initiation | 11/3222 | 17/11632 | **0.043** | 0.039 |
| GO:0010564 | regulation of cell cycle process | 18/3222 | 34/11632 | **0.043** | 0.039 |
| GO:0007015 | actin filament organization | 19/3222 | 37/11632 | **0.048** | 0.044 |
| GO:0051493 | regulation of cytoskeleton organization | 13/3222 | 22/11632 | **0.048** | 0.044 |
| GO:0098542 | defense response to other organism | 15/3222 | 27/11632 | **0.049** | 0.044 |
| **C) Induced in Temperature** | | | | | |
| **ID** | **Description** | **Gene Ratio** | **Term Ratio** | **p** | **q** |
| GO:0007266 | Rho protein signal transduction | 16/1979 | 27/11632 | **< 0.001** | 0 |
| GO:0035556 | intracellular signal transduction | 44/1979 | 143/11632 | **0.008** | 0.008 |
| GO:0006355 | regulation of DNA-templated transcription | 88/1979 | 348/11632 | **0.008** | 0.008 |
| GO:0043161 | proteasome-mediated ubiquitin-dependent protein catabolic process | 30/1979 | 88/11632 | **0.008** | 0.008 |
| GO:0006886 | intracellular protein transport | 40/1979 | 130/11632 | **0.008** | 0.008 |
| GO:0006357 | regulation of transcription by RNA polymerase II | 96/1979 | 395/11632 | **0.009** | 0.009 |
| GO:0006888 | endoplasmic reticulum to Golgi vesicle-mediated transport | 21/1979 | 56/11632 | **0.014** | 0.014 |
| GO:0030968 | endoplasmic reticulum unfolded protein response | 9/1979 | 16/11632 | **0.027** | 0.027 |
| GO:0030433 | ubiquitin-dependent ERAD pathway | 14/1979 | 33/11632 | **0.03** | 0.029 |
| GO:0070534 | protein K63-linked ubiquitination | 15/1979 | 38/11632 | **0.043** | 0.042 |
| GO:0006413 | translational initiation | 11/1979 | 24/11632 | **0.045** | 0.044 |
| GO:0006890 | retrograde vesicle-mediated transport, Golgi to endoplasmic reticulum | 10/1979 | 21/11632 | **0.049** | 0.048 |
| **D) Repressed in Temperature** | | | | | |
| **ID** | **Description** | **Gene Ratio** | **Term Ratio** | **p** | **q** |
| GO:0006412 | translation | 58/2698 | 126/11632 | **< 0.001** | 0 |
| GO:0006260 | DNA replication | 26/2698 | 42/11632 | **< 0.001** | 0 |
| GO:0032508 | DNA duplex unwinding | 17/2698 | 22/11632 | **< 0.001** | 0 |
| GO:0006281 | DNA repair | 42/2698 | 97/11632 | **0.001** | 0.001 |
| GO:0007099 | centriole replication | 9/2698 | 10/11632 | **0.002** | 0.002 |
| GO:0006508 | proteolysis | 96/2698 | 291/11632 | **0.007** | 0.006 |
| GO:0055085 | transmembrane transport | 126/2698 | 403/11632 | **0.007** | 0.007 |
| GO:1902476 | chloride transmembrane transport | 16/2698 | 29/11632 | **0.013** | 0.012 |
| GO:0032970 | regulation of actin filament-based process | 12/2698 | 19/11632 | **0.013** | 0.012 |
| GO:0006302 | double-strand break repair | 16/2698 | 30/11632 | **0.017** | 0.016 |
| GO:0060341 | regulation of cellular localization | 16/2698 | 31/11632 | **0.024** | 0.022 |
| GO:0098655 | monoatomic cation transmembrane transport | 9/2698 | 13/11632 | **0.024** | 0.022 |
| GO:0000727 | double-strand break repair via break-induced replication | 8/2698 | 11/11632 | **0.028** | 0.026 |
| GO:0050896 | response to stimulus | 94/2698 | 301/11632 | **0.028** | 0.026 |
| GO:0070588 | calcium ion transmembrane transport | 22/2698 | 51/11632 | **0.044** | 0.041 |
| GO:0051493 | regulation of cytoskeleton organization | 12/2698 | 22/11632 | **0.048** | 0.044 |
| **E) Induced in Nutrients** | | | | | |
| **ID** | **Description** | **Gene Ratio** | **Term Ratio** | **p** | **q** |
| GO:0030433 | ubiquitin-dependent ERAD pathway | 22/2349 | 33/11632 | **< 0.001** | 0 |
| GO:0043161 | proteasome-mediated ubiquitin-dependent protein catabolic process | 39/2349 | 88/11632 | **< 0.001** | 0 |
| GO:0006511 | ubiquitin-dependent protein catabolic process | 38/2349 | 89/11632 | **< 0.001** | 0 |
| GO:0007266 | Rho protein signal transduction | 17/2349 | 27/11632 | **< 0.001** | 0 |
| GO:0035556 | intracellular signal transduction | 53/2349 | 143/11632 | **< 0.001** | 0 |
| GO:0006886 | intracellular protein transport | 49/2349 | 130/11632 | **< 0.001** | 0 |
| GO:0006355 | regulation of DNA-templated transcription | 103/2349 | 348/11632 | **0.001** | 0.001 |
| GO:0030968 | endoplasmic reticulum unfolded protein response | 11/2349 | 16/11632 | **0.002** | 0.002 |
| GO:0006906 | vesicle fusion | 13/2349 | 21/11632 | **0.002** | 0.002 |
| GO:0006357 | regulation of transcription by RNA polymerase II | 110/2349 | 395/11632 | **0.007** | 0.006 |
| GO:0070534 | protein K63-linked ubiquitination | 18/2349 | 38/11632 | **0.007** | 0.007 |
| GO:0016567 | protein ubiquitination | 41/2349 | 123/11632 | **0.018** | 0.017 |
| GO:0006888 | endoplasmic reticulum to Golgi vesicle-mediated transport | 22/2349 | 56/11632 | **0.032** | 0.03 |
| GO:0006468 | protein phosphorylation | 101/2349 | 375/11632 | **0.032** | 0.03 |
| GO:0006890 | retrograde vesicle-mediated transport, Golgi to endoplasmic reticulum | 11/2349 | 21/11632 | **0.036** | 0.035 |
| **F) Repressed in Nutrients** | | | | | |
| **ID** | **Description** | **Gene Ratio** | **Term Ratio** | **p** | **q** |
| GO:0006412 | translation | 68/3229 | 126/11632 | **< 0.001** | 0 |
| GO:0006260 | DNA replication | 29/3229 | 42/11632 | **< 0.001** | 0 |
| GO:0032508 | DNA duplex unwinding | 17/3229 | 22/11632 | **< 0.001** | 0 |
| GO:0006281 | DNA repair | 47/3229 | 97/11632 | **0.002** | 0.001 |
| GO:0006508 | proteolysis | 113/3229 | 291/11632 | **0.002** | 0.002 |
| GO:0055085 | transmembrane transport | 146/3229 | 403/11632 | **0.009** | 0.009 |
| GO:0002682 | regulation of immune system process | 31/3229 | 63/11632 | **0.017** | 0.016 |
| GO:0032970 | regulation of actin filament-based process | 13/3229 | 19/11632 | **0.017** | 0.016 |
| GO:0005975 | carbohydrate metabolic process | 55/3229 | 131/11632 | **0.017** | 0.016 |
| GO:0030198 | extracellular matrix organization | 23/3229 | 43/11632 | **0.017** | 0.016 |
| GO:1902476 | chloride transmembrane transport | 17/3229 | 29/11632 | **0.023** | 0.022 |
| GO:0060415 | muscle tissue morphogenesis | 8/3229 | 10/11632 | **0.04** | 0.038 |
| GO:0000278 | mitotic cell cycle | 22/3229 | 43/11632 | **0.04** | 0.038 |
| GO:0050896 | response to stimulus | 108/3229 | 301/11632 | **0.044** | 0.041 |
| **G) Induced in Combined** | | | | | |
| **ID** | **Description** | **Gene Ratio** | **Term Ratio** | **p** | **q** |
| GO:0030433 | ubiquitin-dependent ERAD pathway | 20/2149 | 33/11632 | **< 0.001** | 0 |
| GO:0006355 | regulation of DNA-templated transcription | 100/2149 | 348/11632 | **< 0.001** | 0 |
| GO:0035556 | intracellular signal transduction | 48/2149 | 143/11632 | **0.002** | 0.002 |
| GO:0007266 | Rho protein signal transduction | 15/2149 | 27/11632 | **0.002** | 0.002 |
| GO:0043161 | proteasome-mediated ubiquitin-dependent protein catabolic process | 33/2149 | 88/11632 | **0.002** | 0.002 |
| GO:0006357 | regulation of transcription by RNA polymerase II | 106/2149 | 395/11632 | **0.002** | 0.002 |
| GO:0070534 | protein K63-linked ubiquitination | 18/2149 | 38/11632 | **0.003** | 0.003 |
| GO:0006511 | ubiquitin-dependent protein catabolic process | 31/2149 | 89/11632 | **0.011** | 0.011 |
| GO:0006888 | endoplasmic reticulum to Golgi vesicle-mediated transport | 22/2149 | 56/11632 | **0.012** | 0.012 |
| GO:0060294 | cilium movement involved in cell motility | 8/2149 | 12/11632 | **0.017** | 0.016 |
| GO:0007018 | microtubule-based movement | 30/2149 | 90/11632 | **0.025** | 0.024 |
| GO:0030968 | endoplasmic reticulum unfolded protein response | 9/2149 | 16/11632 | **0.035** | 0.034 |
| GO:0006886 | intracellular protein transport | 39/2149 | 130/11632 | **0.037** | 0.036 |
| GO:0006468 | protein phosphorylation | 93/2149 | 375/11632 | **0.044** | 0.042 |
| GO:0003341 | cilium movement | 11/2149 | 23/11632 | **0.044** | 0.042 |
| GO:0033209 | tumor necrosis factor-mediated signaling pathway | 15/2149 | 37/11632 | **0.047** | 0.046 |
| **H) Repressed in Combined** | | | | | |
| **ID** | **Description** | **Gene Ratio** | **Term Ratio** | **p** | **q** |
| GO:0006260 | DNA replication | 31/3106 | 42/11632 | **< 0.001** | 0 |
| GO:0006281 | DNA repair | 53/3106 | 97/11632 | **< 0.001** | 0 |
| GO:0006412 | translation | 61/3106 | 126/11632 | **< 0.001** | 0 |
| GO:0032508 | DNA duplex unwinding | 17/3106 | 22/11632 | **< 0.001** | 0 |
| GO:0000727 | double-strand break repair via break-induced replication | 10/3106 | 11/11632 | **0.002** | 0.002 |
| GO:0055085 | transmembrane transport | 143/3106 | 403/11632 | **0.004** | 0.004 |
| GO:0098655 | monoatomic cation transmembrane transport | 10/3106 | 13/11632 | **0.016** | 0.015 |
| GO:1902476 | chloride transmembrane transport | 17/3106 | 29/11632 | **0.016** | 0.015 |
| GO:0002682 | regulation of immune system process | 30/3106 | 63/11632 | **0.016** | 0.015 |
| GO:0006635 | fatty acid beta-oxidation | 14/3106 | 22/11632 | **0.016** | 0.015 |
| GO:0000723 | telomere maintenance | 13/3106 | 20/11632 | **0.018** | 0.017 |
| GO:0006298 | mismatch repair | 11/3106 | 16/11632 | **0.024** | 0.022 |
| GO:0002376 | immune system process | 42/3106 | 100/11632 | **0.026** | 0.024 |
| GO:0007156 | homophilic cell adhesion via plasma membrane adhesion molecules | 31/3106 | 69/11632 | **0.031** | 0.028 |
| GO:0006508 | proteolysis | 102/3106 | 291/11632 | **0.031** | 0.028 |
| GO:0032970 | regulation of actin filament-based process | 12/3106 | 19/11632 | **0.031** | 0.028 |
| GO:0006261 | DNA-templated DNA replication | 10/3106 | 15/11632 | **0.044** | 0.04 |

**Table S9. Results of Gene Ontology (GO) functional enrichment analysis throughout thermal challenge.** Result of overrepresentation analysis of gene ontology (GO) terms of biological processes for genes up- and down-regulated in temperature-primed **(A, C, E, G, I, K**) and naive (**B, D, F, H, J, L**) corals at each time point (C1: **A-B**, C2: **C-D**, C3: **E-F**, C4: **G-H**, C5: **I-J**, C6: **K-L**) throughout acute heating during the thermal challenge (**Figure S9**). For each GO term, the gene ratio (significant genes / total genes in term), term ratio (total genes in term / total genes), p value (bold: p < 0.05), and q value are listed.

| **A) C1: Temperature-Primed** | | | | | | |
| --- | --- | --- | --- | --- | --- | --- |
| **Regulation** | **ID** | **Description** | **Gene Ratio** | **Term Ratio** | **p** | **q** |
| Induced | GO:0061024 | membrane organization | 2/19 | 26/13081 | **0.029** | 0.019 |
| Repressed | GO:0006355 | regulation of DNA-templated transcription | 10/87 | 386/13081 | **0.026** | 0.022 |
| **B) C1: Naive** | | | | | | |
| **Regulation** | **ID** | **Description** | **Gene Ratio** | **Term Ratio** | **p** | **q** |
| Repressed | GO:0006355 | regulation of DNA-templated transcription | 17/206 | 386/13081 | **0.016** | 0.015 |
| Repressed | GO:0006357 | regulation of transcription by RNA polymerase II | 18/206 | 430/13081 | **0.016** | 0.015 |
| Repressed | GO:0035556 | intracellular signal transduction | 9/206 | 150/13081 | **0.041** | 0.038 |
| **C) C2: Temperature-Primed** | | | | | | |
| **Regulation** | **ID** | **Description** | **Gene Ratio** | **Term Ratio** | **p** | **q** |
| Induced | GO:0006457 | protein folding | 12/319 | 78/13081 | **< 0.001** | 0 |
| Induced | GO:0016567 | protein ubiquitination | 13/319 | 131/13081 | **0.003** | 0.003 |
| Induced | GO:0034976 | response to endoplasmic reticulum stress | 4/319 | 11/13081 | **0.01** | 0.01 |
| Induced | GO:0042981 | regulation of apoptotic process | 10/319 | 116/13081 | **0.042** | 0.039 |
| **D) C2: Naive** | | | | | | |
| **Regulation** | **ID** | **Description** | **Gene Ratio** | **Term Ratio** | **p** | **q** |
| Induced | GO:0006457 | protein folding | 12/329 | 78/13081 | **< 0.001** | 0 |
| Induced | GO:0006357 | regulation of transcription by RNA polymerase II | 26/329 | 430/13081 | **0.005** | 0.005 |
| Induced | GO:0034976 | response to endoplasmic reticulum stress | 4/329 | 11/13081 | **0.011** | 0.011 |
| Induced | GO:0030433 | ubiquitin-dependent ERAD pathway | 6/329 | 33/13081 | **0.011** | 0.011 |
| Induced | GO:0016567 | protein ubiquitination | 11/329 | 131/13081 | **0.028** | 0.026 |
| Induced | GO:0000226 | microtubule cytoskeleton organization | 7/329 | 59/13081 | **0.033** | 0.03 |
| Induced | GO:0006355 | regulation of DNA-templated transcription | 21/329 | 386/13081 | **0.033** | 0.03 |
| Induced | GO:0007017 | microtubule-based process | 6/329 | 45/13081 | **0.033** | 0.03 |
| **E) C3: Temperature-Primed** | | | | | | |
| **Regulation** | **ID** | **Description** | **Gene Ratio** | **Term Ratio** | **p** | **q** |
| Induced | GO:0006457 | protein folding | 18/723 | 78/13081 | **< 0.001** | 0 |
| Induced | GO:0007266 | Rho protein signal transduction | 9/723 | 28/13081 | **0.002** | 0.002 |
| Induced | GO:0006357 | regulation of transcription by RNA polymerase II | 46/723 | 430/13081 | **0.002** | 0.002 |
| Induced | GO:0016567 | protein ubiquitination | 19/723 | 131/13081 | **0.011** | 0.011 |
| Induced | GO:0016477 | cell migration | 10/723 | 45/13081 | **0.012** | 0.011 |
| Induced | GO:0034976 | response to endoplasmic reticulum stress | 5/723 | 11/13081 | **0.013** | 0.012 |
| Induced | GO:0042981 | regulation of apoptotic process | 17/723 | 116/13081 | **0.013** | 0.012 |
| Induced | GO:0032970 | regulation of actin filament-based process | 6/723 | 20/13081 | **0.03** | 0.029 |
| **F) C3: Naive** | | | | | | |
| **Regulation** | **ID** | **Description** | **Gene Ratio** | **Term Ratio** | **p** | **q** |
| Induced | GO:0007266 | Rho protein signal transduction | 12/912 | 28/13081 | **< 0.001** | 0 |
| Induced | GO:0016567 | protein ubiquitination | 24/912 | 131/13081 | **0.002** | 0.002 |
| Induced | GO:0006355 | regulation of DNA-templated transcription | 50/912 | 386/13081 | **0.002** | 0.002 |
| Induced | GO:0006357 | regulation of transcription by RNA polymerase II | 54/912 | 430/13081 | **0.002** | 0.002 |
| Induced | GO:0016477 | cell migration | 12/912 | 45/13081 | **0.004** | 0.004 |
| Induced | GO:0006457 | protein folding | 15/912 | 78/13081 | **0.019** | 0.018 |
| Induced | GO:0030433 | ubiquitin-dependent ERAD pathway | 9/912 | 33/13081 | **0.019** | 0.018 |
| Induced | GO:0007163 | establishment or maintenance of cell polarity | 6/912 | 15/13081 | **0.019** | 0.018 |
| Induced | GO:0033209 | tumor necrosis factor-mediated signaling pathway | 10/912 | 41/13081 | **0.02** | 0.019 |
| Induced | GO:0070534 | protein K63-linked ubiquitination | 10/912 | 42/13081 | **0.022** | 0.021 |
| Induced | GO:0034976 | response to endoplasmic reticulum stress | 5/912 | 11/13081 | **0.022** | 0.021 |
| Induced | GO:0030162 | regulation of proteolysis | 5/912 | 12/13081 | **0.033** | 0.031 |
| Induced | GO:0042981 | regulation of apoptotic process | 18/912 | 116/13081 | **0.038** | 0.036 |
| **G) C4: Temperature-Primed** | | | | | | |
| **Regulation** | **ID** | **Description** | **Gene Ratio** | **Term Ratio** | **p** | **q** |
| Induced | GO:0006355 | regulation of DNA-templated transcription | 102/1963 | 386/13081 | **< 0.001** | 0 |
| Induced | GO:0006357 | regulation of transcription by RNA polymerase II | 104/1963 | 430/13081 | **< 0.001** | 0 |
| Induced | GO:0007266 | Rho protein signal transduction | 16/1963 | 28/13081 | **< 0.001** | 0 |
| Induced | GO:0016567 | protein ubiquitination | 40/1963 | 131/13081 | **0.001** | 0.001 |
| Induced | GO:0006511 | ubiquitin-dependent protein catabolic process | 30/1963 | 92/13081 | **0.002** | 0.002 |
| Induced | GO:0006468 | protein phosphorylation | 90/1963 | 407/13081 | **0.006** | 0.006 |
| Induced | GO:0035556 | intracellular signal transduction | 40/1963 | 150/13081 | **0.011** | 0.011 |
| Induced | GO:0043161 | proteasome-mediated ubiquitin-dependent protein catabolic process | 27/1963 | 92/13081 | **0.022** | 0.021 |
| Induced | GO:0007034 | vacuolar transport | 8/1963 | 15/13081 | **0.034** | 0.032 |
| Induced | GO:0032436 | positive regulation of proteasomal ubiquitin-dependent protein catabolic process | 10/1963 | 22/13081 | **0.034** | 0.032 |
| Induced | GO:0006888 | endoplasmic reticulum to Golgi vesicle-mediated transport | 19/1963 | 59/13081 | **0.034** | 0.032 |
| Repressed | GO:0006412 | translation | 72/3913 | 132/13081 | **< 0.001** | 0 |
| Repressed | GO:0006260 | DNA replication | 34/3913 | 50/13081 | **< 0.001** | 0 |
| Repressed | GO:0051225 | spindle assembly | 13/3913 | 14/13081 | **< 0.001** | 0 |
| Repressed | GO:0006281 | DNA repair | 56/3913 | 113/13081 | **0.001** | 0.001 |
| Repressed | GO:0000727 | double-strand break repair via break-induced replication | 11/3913 | 12/13081 | **0.002** | 0.002 |
| Repressed | GO:0000712 | resolution of meiotic recombination intermediates | 10/3913 | 11/13081 | **0.005** | 0.004 |
| Repressed | GO:0007099 | centriole replication | 9/3913 | 10/13081 | **0.012** | 0.011 |
| Repressed | GO:0006635 | fatty acid beta-oxidation | 15/3913 | 22/13081 | **0.017** | 0.017 |
| Repressed | GO:0000278 | mitotic cell cycle | 24/3913 | 43/13081 | **0.023** | 0.022 |
| Repressed | GO:0006302 | double-strand break repair | 19/3913 | 32/13081 | **0.03** | 0.029 |
| **H) C4: Naive** | | | | | | |
| **Regulation** | **ID** | **Description** | **Gene Ratio** | **Term Ratio** | **p** | **q** |
| Induced | GO:0006355 | regulation of DNA-templated transcription | 102/2051 | 386/13081 | **< 0.001** | 0 |
| Induced | GO:0006357 | regulation of transcription by RNA polymerase II | 104/2051 | 430/13081 | **< 0.001** | 0 |
| Induced | GO:0016567 | protein ubiquitination | 42/2051 | 131/13081 | **< 0.001** | 0 |
| Induced | GO:0007266 | Rho protein signal transduction | 15/2051 | 28/13081 | **0.001** | 0.001 |
| Induced | GO:0006511 | ubiquitin-dependent protein catabolic process | 32/2051 | 92/13081 | **0.001** | 0.001 |
| Induced | GO:0035556 | intracellular signal transduction | 44/2051 | 150/13081 | **0.001** | 0.001 |
| Induced | GO:0006468 | protein phosphorylation | 93/2051 | 407/13081 | **0.006** | 0.006 |
| Induced | GO:0043161 | proteasome-mediated ubiquitin-dependent protein catabolic process | 29/2051 | 92/13081 | **0.007** | 0.007 |
| Induced | GO:0006886 | intracellular protein transport | 38/2051 | 135/13081 | **0.009** | 0.009 |
| Induced | GO:0030433 | ubiquitin-dependent ERAD pathway | 14/2051 | 33/13081 | **0.012** | 0.012 |
| Induced | GO:0006888 | endoplasmic reticulum to Golgi vesicle-mediated transport | 20/2051 | 59/13081 | **0.019** | 0.019 |
| Induced | GO:0034976 | response to endoplasmic reticulum stress | 7/2051 | 11/13081 | **0.019** | 0.019 |
| Induced | GO:0032436 | positive regulation of proteasomal ubiquitin-dependent protein catabolic process | 10/2051 | 22/13081 | **0.039** | 0.037 |
| Repressed | GO:0006412 | translation | 72/3976 | 132/13081 | **< 0.001** | 0 |
| Repressed | GO:0006260 | DNA replication | 33/3976 | 50/13081 | **< 0.001** | 0 |
| Repressed | GO:0006281 | DNA repair | 58/3976 | 113/13081 | **< 0.001** | 0 |
| Repressed | GO:0006284 | base-excision repair | 13/3976 | 16/13081 | **0.006** | 0.006 |
| Repressed | GO:0007099 | centriole replication | 9/3976 | 10/13081 | **0.017** | 0.016 |
| Repressed | GO:0006302 | double-strand break repair | 20/3976 | 32/13081 | **0.017** | 0.016 |
| Repressed | GO:0000727 | double-strand break repair via break-induced replication | 10/3976 | 12/13081 | **0.02** | 0.019 |
| Repressed | GO:0006635 | fatty acid beta-oxidation | 15/3976 | 22/13081 | **0.021** | 0.021 |
| Repressed | GO:0000712 | resolution of meiotic recombination intermediates | 9/3976 | 11/13081 | **0.043** | 0.041 |
| **I) C5: Temperature-Primed** | | | | | | |
| **Regulation** | **ID** | **Description** | **Gene Ratio** | **Term Ratio** | **p** | **q** |
| Induced | GO:0007266 | Rho protein signal transduction | 15/1782 | 28/13081 | **< 0.001** | 0 |
| Induced | GO:0016567 | protein ubiquitination | 35/1782 | 131/13081 | **0.008** | 0.008 |
| Induced | GO:0006888 | endoplasmic reticulum to Golgi vesicle-mediated transport | 20/1782 | 59/13081 | **0.008** | 0.008 |
| Induced | GO:0006355 | regulation of DNA-templated transcription | 80/1782 | 386/13081 | **0.008** | 0.008 |
| Induced | GO:0006511 | ubiquitin-dependent protein catabolic process | 26/1782 | 92/13081 | **0.015** | 0.015 |
| Induced | GO:0034976 | response to endoplasmic reticulum stress | 7/1782 | 11/13081 | **0.015** | 0.015 |
| Induced | GO:0030433 | ubiquitin-dependent ERAD pathway | 13/1782 | 33/13081 | **0.016** | 0.016 |
| Induced | GO:0006886 | intracellular protein transport | 33/1782 | 135/13081 | **0.032** | 0.031 |
| Induced | GO:0006457 | protein folding | 22/1782 | 78/13081 | **0.032** | 0.031 |
| Induced | GO:0006357 | regulation of transcription by RNA polymerase II | 82/1782 | 430/13081 | **0.045** | 0.044 |
| Repressed | GO:0006260 | DNA replication | 33/3779 | 50/13081 | **< 0.001** | 0 |
| Repressed | GO:0006281 | DNA repair | 57/3779 | 113/13081 | **< 0.001** | 0 |
| Repressed | GO:0006270 | DNA replication initiation | 13/3779 | 17/13081 | **0.013** | 0.013 |
| Repressed | GO:0007099 | centriole replication | 9/3779 | 10/13081 | **0.015** | 0.015 |
| Repressed | GO:0000727 | double-strand break repair via break-induced replication | 10/3779 | 12/13081 | **0.015** | 0.015 |
| Repressed | GO:0006635 | fatty acid beta-oxidation | 15/3779 | 22/13081 | **0.015** | 0.015 |
| Repressed | GO:0055085 | transmembrane transport | 164/3779 | 451/13081 | **0.025** | 0.024 |
| Repressed | GO:0000723 | telomere maintenance | 14/3779 | 21/13081 | **0.027** | 0.026 |
| Repressed | GO:0009653 | anatomical structure morphogenesis | 43/3779 | 95/13081 | **0.033** | 0.032 |
| Repressed | GO:0006508 | proteolysis | 119/3779 | 319/13081 | **0.038** | 0.037 |
| **J) C5: Naive** | | | | | | |
| **Regulation** | **ID** | **Description** | **Gene Ratio** | **Term Ratio** | **p** | **q** |
| Induced | GO:0007266 | Rho protein signal transduction | 16/1878 | 28/13081 | **< 0.001** | 0 |
| Induced | GO:0006355 | regulation of DNA-templated transcription | 91/1878 | 386/13081 | **< 0.001** | 0 |
| Induced | GO:0016567 | protein ubiquitination | 40/1878 | 131/13081 | **< 0.001** | 0 |
| Induced | GO:0006511 | ubiquitin-dependent protein catabolic process | 31/1878 | 92/13081 | **< 0.001** | 0 |
| Induced | GO:0070534 | protein K63-linked ubiquitination | 17/1878 | 42/13081 | **0.003** | 0.003 |
| Induced | GO:0006357 | regulation of transcription by RNA polymerase II | 92/1878 | 430/13081 | **0.003** | 0.003 |
| Induced | GO:0033209 | tumor necrosis factor-mediated signaling pathway | 16/1878 | 41/13081 | **0.007** | 0.007 |
| Induced | GO:0006888 | endoplasmic reticulum to Golgi vesicle-mediated transport | 20/1878 | 59/13081 | **0.009** | 0.008 |
| Induced | GO:0043161 | proteasome-mediated ubiquitin-dependent protein catabolic process | 27/1878 | 92/13081 | **0.009** | 0.009 |
| Induced | GO:0034976 | response to endoplasmic reticulum stress | 7/1878 | 11/13081 | **0.013** | 0.012 |
| Induced | GO:0006886 | intracellular protein transport | 35/1878 | 135/13081 | **0.014** | 0.013 |
| Induced | GO:0030433 | ubiquitin-dependent ERAD pathway | 13/1878 | 33/13081 | **0.016** | 0.016 |
| Induced | GO:0032436 | positive regulation of proteasomal ubiquitin-dependent protein catabolic process | 10/1878 | 22/13081 | **0.019** | 0.018 |
| Induced | GO:0035556 | intracellular signal transduction | 37/1878 | 150/13081 | **0.021** | 0.02 |
| Induced | GO:0006468 | protein phosphorylation | 82/1878 | 407/13081 | **0.027** | 0.026 |
| Induced | GO:0006457 | protein folding | 22/1878 | 78/13081 | **0.037** | 0.035 |
| Repressed | GO:0006412 | translation | 67/3636 | 132/13081 | **< 0.001** | 0 |
| Repressed | GO:0006260 | DNA replication | 33/3636 | 50/13081 | **< 0.001** | 0 |
| Repressed | GO:0006281 | DNA repair | 53/3636 | 113/13081 | **0.002** | 0.002 |
| Repressed | GO:0007099 | centriole replication | 9/3636 | 10/13081 | **0.011** | 0.01 |
| Repressed | GO:1902476 | chloride transmembrane transport | 20/3636 | 34/13081 | **0.017** | 0.016 |
| Repressed | GO:0055085 | transmembrane transport | 159/3636 | 451/13081 | **0.02** | 0.019 |
| Repressed | GO:0006508 | proteolysis | 117/3636 | 319/13081 | **0.02** | 0.019 |
| Repressed | GO:0006270 | DNA replication initiation | 12/3636 | 17/13081 | **0.02** | 0.019 |
| Repressed | GO:0002684 | positive regulation of immune system process | 9/3636 | 11/13081 | **0.02** | 0.019 |
| Repressed | GO:0030198 | extracellular matrix organization | 24/3636 | 46/13081 | **0.024** | 0.022 |
| Repressed | GO:0006635 | fatty acid beta-oxidation | 14/3636 | 22/13081 | **0.026** | 0.024 |
| Repressed | GO:0006298 | mismatch repair | 12/3636 | 18/13081 | **0.033** | 0.03 |
| Repressed | GO:0000727 | double-strand break repair via break-induced replication | 9/3636 | 12/13081 | **0.042** | 0.039 |
| Repressed | GO:0000723 | telomere maintenance | 13/3636 | 21/13081 | **0.048** | 0.044 |
| **K) C6: Temperature-Primed** | | | | | | |
| **Regulation** | **ID** | **Description** | **Gene Ratio** | **Term Ratio** | **p** | **q** |
| Induced | GO:0007266 | Rho protein signal transduction | 15/1436 | 28/13081 | **< 0.001** | 0 |
| Induced | GO:0006355 | regulation of DNA-templated transcription | 72/1436 | 386/13081 | **0.001** | 0.001 |
| Induced | GO:0035556 | intracellular signal transduction | 34/1436 | 150/13081 | **0.004** | 0.004 |
| Induced | GO:0006357 | regulation of transcription by RNA polymerase II | 75/1436 | 430/13081 | **0.004** | 0.004 |
| Induced | GO:0006413 | translational initiation | 10/1436 | 24/13081 | **0.012** | 0.011 |
| Induced | GO:0030433 | ubiquitin-dependent ERAD pathway | 11/1436 | 33/13081 | **0.045** | 0.044 |
| Repressed | GO:0006260 | DNA replication | 29/2600 | 50/13081 | **< 0.001** | 0 |
| Repressed | GO:0006412 | translation | 54/2600 | 132/13081 | **< 0.001** | 0 |
| Repressed | GO:0006281 | DNA repair | 43/2600 | 113/13081 | **0.001** | 0.001 |
| Repressed | GO:0006508 | proteolysis | 93/2600 | 319/13081 | **0.005** | 0.005 |
| Repressed | GO:0055085 | transmembrane transport | 124/2600 | 451/13081 | **0.005** | 0.005 |
| Repressed | GO:0098655 | monoatomic cation transmembrane transport | 10/2600 | 15/13081 | **0.009** | 0.008 |
| Repressed | GO:0032508 | DNA duplex unwinding | 17/2600 | 35/13081 | **0.009** | 0.008 |
| Repressed | GO:0006302 | double-strand break repair | 16/2600 | 32/13081 | **0.009** | 0.008 |
| Repressed | GO:0032970 | regulation of actin filament-based process | 11/2600 | 20/13081 | **0.032** | 0.029 |
| Repressed | GO:0000727 | double-strand break repair via break-induced replication | 8/2600 | 12/13081 | **0.032** | 0.029 |
| Repressed | GO:0060415 | muscle tissue morphogenesis | 7/2600 | 10/13081 | **0.043** | 0.039 |
| **L) C6: Naive** | | | | | | |
| **Regulation** | **ID** | **Description** | **Gene Ratio** | **Term Ratio** | **p** | **q** |
| Induced | GO:0043161 | proteasome-mediated ubiquitin-dependent protein catabolic process | 32/1689 | 92/13081 | **< 0.001** | 0 |
| Induced | GO:0006355 | regulation of DNA-templated transcription | 85/1689 | 386/13081 | **< 0.001** | 0 |
| Induced | GO:0007266 | Rho protein signal transduction | 14/1689 | 28/13081 | **< 0.001** | 0 |
| Induced | GO:0006357 | regulation of transcription by RNA polymerase II | 88/1689 | 430/13081 | **0.001** | 0.001 |
| Induced | GO:0006511 | ubiquitin-dependent protein catabolic process | 27/1689 | 92/13081 | **0.003** | 0.002 |
| Induced | GO:0016567 | protein ubiquitination | 34/1689 | 131/13081 | **0.004** | 0.003 |
| Induced | GO:0010498 | proteasomal protein catabolic process | 8/1689 | 13/13081 | **0.004** | 0.004 |
| Induced | GO:0051603 | proteolysis involved in protein catabolic process | 17/1689 | 50/13081 | **0.007** | 0.007 |
| Induced | GO:0030433 | ubiquitin-dependent ERAD pathway | 13/1689 | 33/13081 | **0.007** | 0.007 |
| Induced | GO:0035556 | intracellular signal transduction | 36/1689 | 150/13081 | **0.007** | 0.007 |
| Induced | GO:0006468 | protein phosphorylation | 77/1689 | 407/13081 | **0.015** | 0.015 |
| Induced | GO:0006413 | translational initiation | 10/1689 | 24/13081 | **0.02** | 0.019 |
| Induced | GO:0006886 | intracellular protein transport | 31/1689 | 135/13081 | **0.036** | 0.035 |
| Induced | GO:0006906 | vesicle fusion | 9/1689 | 22/13081 | **0.038** | 0.037 |
| Repressed | GO:0006260 | DNA replication | 32/3272 | 50/13081 | **< 0.001** | 0 |
| Repressed | GO:0006508 | proteolysis | 120/3272 | 319/13081 | **< 0.001** | 0 |
| Repressed | GO:0006281 | DNA repair | 52/3272 | 113/13081 | **< 0.001** | 0 |
| Repressed | GO:0055085 | transmembrane transport | 155/3272 | 451/13081 | **0.001** | 0.001 |
| Repressed | GO:0006412 | translation | 55/3272 | 132/13081 | **0.002** | 0.002 |
| Repressed | GO:0030198 | extracellular matrix organization | 25/3272 | 46/13081 | **0.002** | 0.002 |
| Repressed | GO:0032970 | regulation of actin filament-based process | 14/3272 | 20/13081 | **0.002** | 0.002 |
| Repressed | GO:0006302 | double-strand break repair | 19/3272 | 32/13081 | **0.002** | 0.002 |
| Repressed | GO:0000727 | double-strand break repair via break-induced replication | 10/3272 | 12/13081 | **0.002** | 0.002 |
| Repressed | GO:0060341 | regulation of cellular localization | 18/3272 | 32/13081 | **0.009** | 0.008 |
| Repressed | GO:0032508 | DNA duplex unwinding | 19/3272 | 35/13081 | **0.011** | 0.009 |
| Repressed | GO:1903566 | positive regulation of protein localization to cilium | 9/3272 | 12/13081 | **0.017** | 0.015 |
| Repressed | GO:0007099 | centriole replication | 8/3272 | 10/13081 | **0.017** | 0.015 |
| Repressed | GO:0060415 | muscle tissue morphogenesis | 8/3272 | 10/13081 | **0.017** | 0.015 |
| Repressed | GO:1902476 | chloride transmembrane transport | 18/3272 | 34/13081 | **0.017** | 0.015 |
| Repressed | GO:0006635 | fatty acid beta-oxidation | 13/3272 | 22/13081 | **0.023** | 0.021 |
| Repressed | GO:0051493 | regulation of cytoskeleton organization | 13/3272 | 22/13081 | **0.023** | 0.021 |
| Repressed | GO:0007015 | actin filament organization | 19/3272 | 38/13081 | **0.023** | 0.021 |
| Repressed | GO:0006261 | DNA-templated DNA replication | 10/3272 | 15/13081 | **0.023** | 0.021 |
| Repressed | GO:0098655 | monoatomic cation transmembrane transport | 10/3272 | 15/13081 | **0.023** | 0.021 |
| Repressed | GO:0050896 | response to stimulus | 115/3272 | 355/13081 | **0.026** | 0.023 |
| Repressed | GO:0000724 | double-strand break repair via homologous recombination | 26/3272 | 60/13081 | **0.039** | 0.035 |
| Repressed | GO:0009103 | lipopolysaccharide biosynthetic process | 14/3272 | 26/13081 | **0.039** | 0.035 |
| Repressed | GO:0000723 | telomere maintenance | 12/3272 | 21/13081 | **0.041** | 0.037 |
| Repressed | GO:0006950 | response to stress | 53/3272 | 147/13081 | **0.042** | 0.038 |

**Table S10. Results of Gene Ontology (GO) functional enrichment analysis in gene modules displaying primed transcriptional resilience during thermal challenge.** Result of overrepresentation analysis of gene ontology (GO) terms of biological processes for genes where expression was stronger in the naive treatment compared to the temperature-primed in **A)** upregulated (ME6, 2, 9) and **B)** downregulated (ME1) WGCNA gene modules (**Figures 4 and S10**). For each GO term, the gene ratio (significant genes / total genes in term), term ratio (total genes in term / total genes), p value (bold: p < 0.05), and q value are listed.

| **A) Upregulated** | | | | | |
| --- | --- | --- | --- | --- | --- |
| **ID** | **Description** | **Gene Ratio** | **Term Ratio** | **p** | **q** |
| GO:0007266 | Rho protein signal transduction | 15/1729 | 25/9225 | **0.002** | 0.002 |
| GO:0006906 | vesicle fusion | 12/1729 | 18/9225 | **0.002** | 0.002 |
| GO:0006355 | regulation of DNA-templated transcription | 79/1729 | 270/9225 | **0.002** | 0.002 |
| GO:0006357 | regulation of transcription by RNA polymerase II | 89/1729 | 313/9225 | **0.002** | 0.002 |
| GO:0033209 | tumor necrosis factor-mediated signaling pathway | 16/1729 | 32/9225 | **0.005** | 0.004 |
| GO:0070534 | protein K63-linked ubiquitination | 16/1729 | 32/9225 | **0.005** | 0.004 |
| GO:0000132 | establishment of mitotic spindle orientation | 14/1729 | 30/9225 | **0.028** | 0.028 |
| GO:0032436 | positive regulation of proteasomal ubiquitin-dependent protein catabolic process | 8/1729 | 13/9225 | **0.042** | 0.042 |
| GO:0006511 | ubiquitin-dependent protein catabolic process | 23/1729 | 64/9225 | **0.042** | 0.042 |
| **B) Downregulated** | | | | | |
| **ID** | **Description** | **Gene Ratio** | **Term Ratio** | **p** | **q** |
| GO:0006281 | DNA repair | 55/3751 | 79/9225 | **< 0.001** | 0 |
| GO:0006260 | DNA replication | 29/3751 | 37/9225 | **0.001** | 0.001 |
| GO:0000723 | telomere maintenance | 14/3751 | 17/9225 | **0.05** | 0.048 |
| GO:0006302 | double-strand break repair | 18/3751 | 24/9225 | **0.05** | 0.048 |
| GO:0032508 | DNA duplex unwinding | 15/3751 | 19/9225 | **0.05** | 0.048 |
| GO:1902476 | chloride transmembrane transport | 19/3751 | 26/9225 | **0.05** | 0.048 |
| GO:0005975 | carbohydrate metabolic process | 61/3751 | 109/9225 | **0.05** | 0.048 |
| GO:0051225 | spindle assembly | 10/3751 | 11/9225 | **0.05** | 0.048 |

**Supplemental Figures**

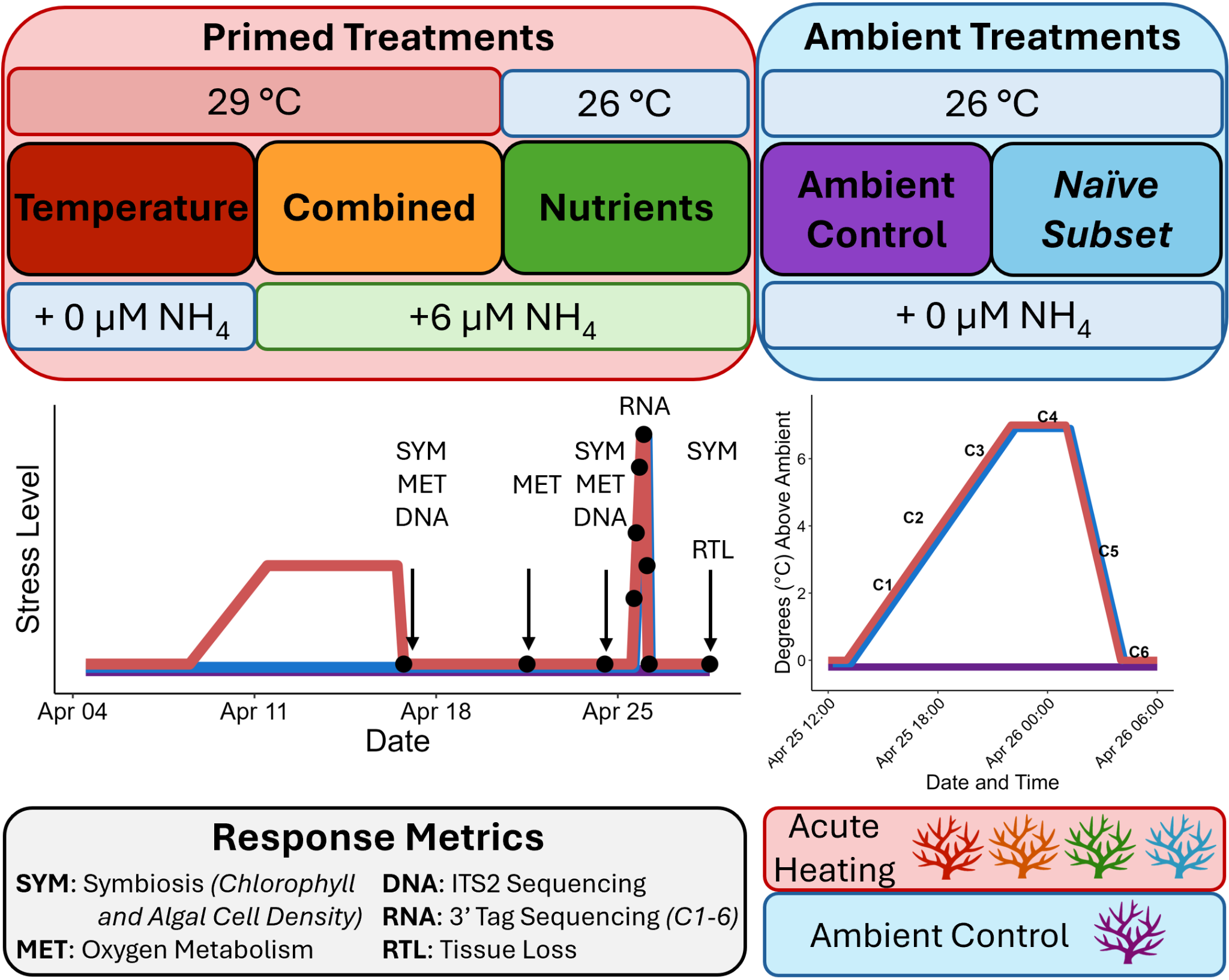

**Figure S1. Experimental design.** Priming treatments, including elevated temperature, nutrients, or the combination of both, compared to ambient conditions. The full experimental time series of priming, recovery, and the acute thermal challenge is shown with sampling time points and the responses (symbiosis, metabolism, symbiont community composition [DNA], gene expression [RNA], and tissue loss) characterized at each. The gene expression sampling time points during acute heating (C1-C6) are shown.

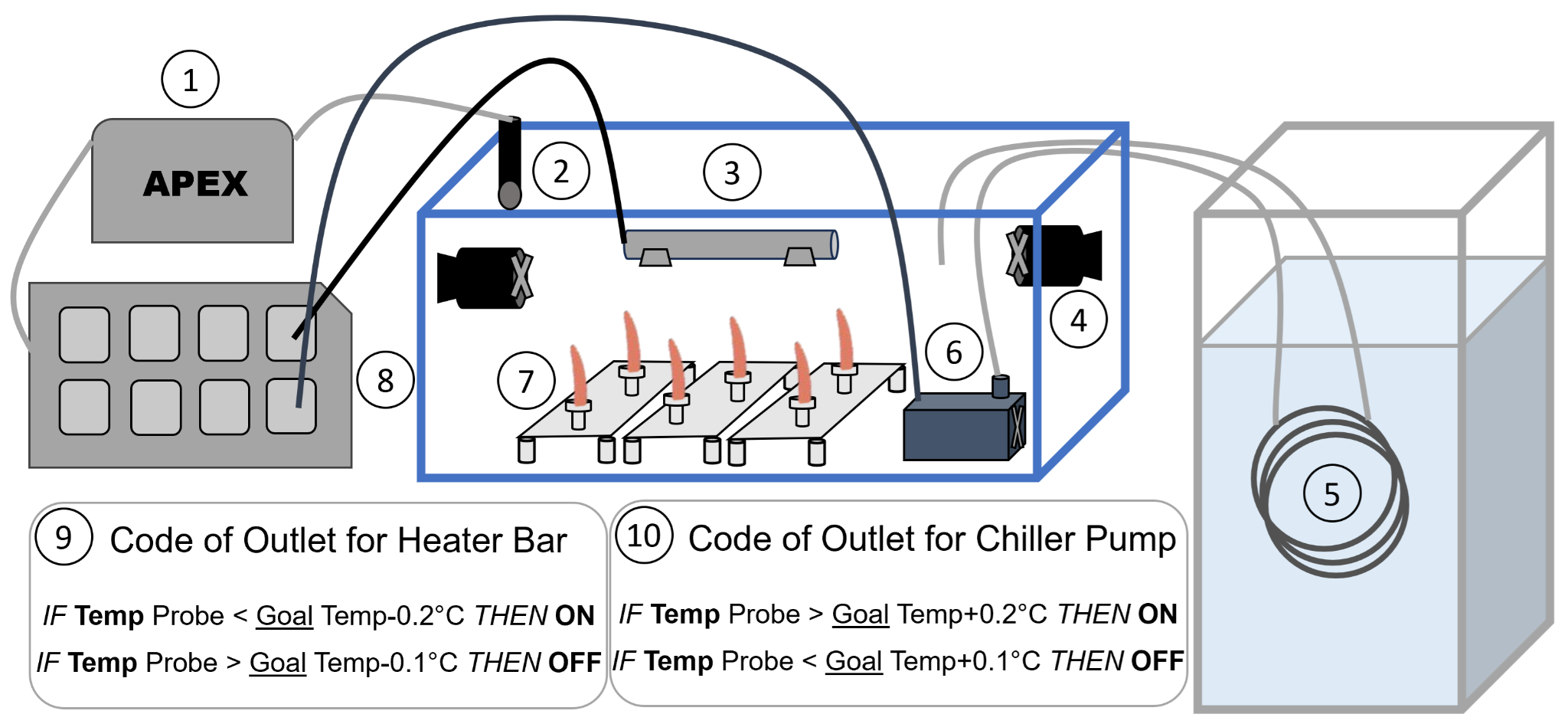

**Figure S2.** **Experimental tank and temperature control design.** In each 150 L tank, temperature was controlled using the Neptune APEX system. Each tank was controlled by an APEX module (1) connected to an APEX temperature probe (2), which was placed on the opposite side of the tank compared to the source of water inflow. A 300 W titanium heater bar (Bulk Reef Supply) (3) was attached to the tank with suction cups, not adjacent to the temperature probe. Two 950 GPH powerheads (Aqueon) (4) were attached to the tank with suction cups, diagonal from each other, to create a circular water flow, including flow across the heater bar. A heat exchange “chiller” system was created with a stainless steel coil within an ice bath (5) attached to aquarium tubing, through which tank water was pumped with a 500 GPH pump (Danner Manufacturing) (6) for heat exchange. The inflow of chilled water was placed near a power head but opposite the temperature probe. Coral fragments were attached to three egg crate grids (7) per tank to allow for rotation of positions within a tank. The heater bar and “chiller” pump were each plugged into specific outlets on an APEX Energy Bar (EB832) (8). The temperature for each tank was controlled by specifying ON and OFF conditions for the outlet of the heater bar (9) and the outlet of the chiller pump (10) based on the temperature measured by the APEX temperature probe, relative to the goal temperature at the specific date and time within the experiment.

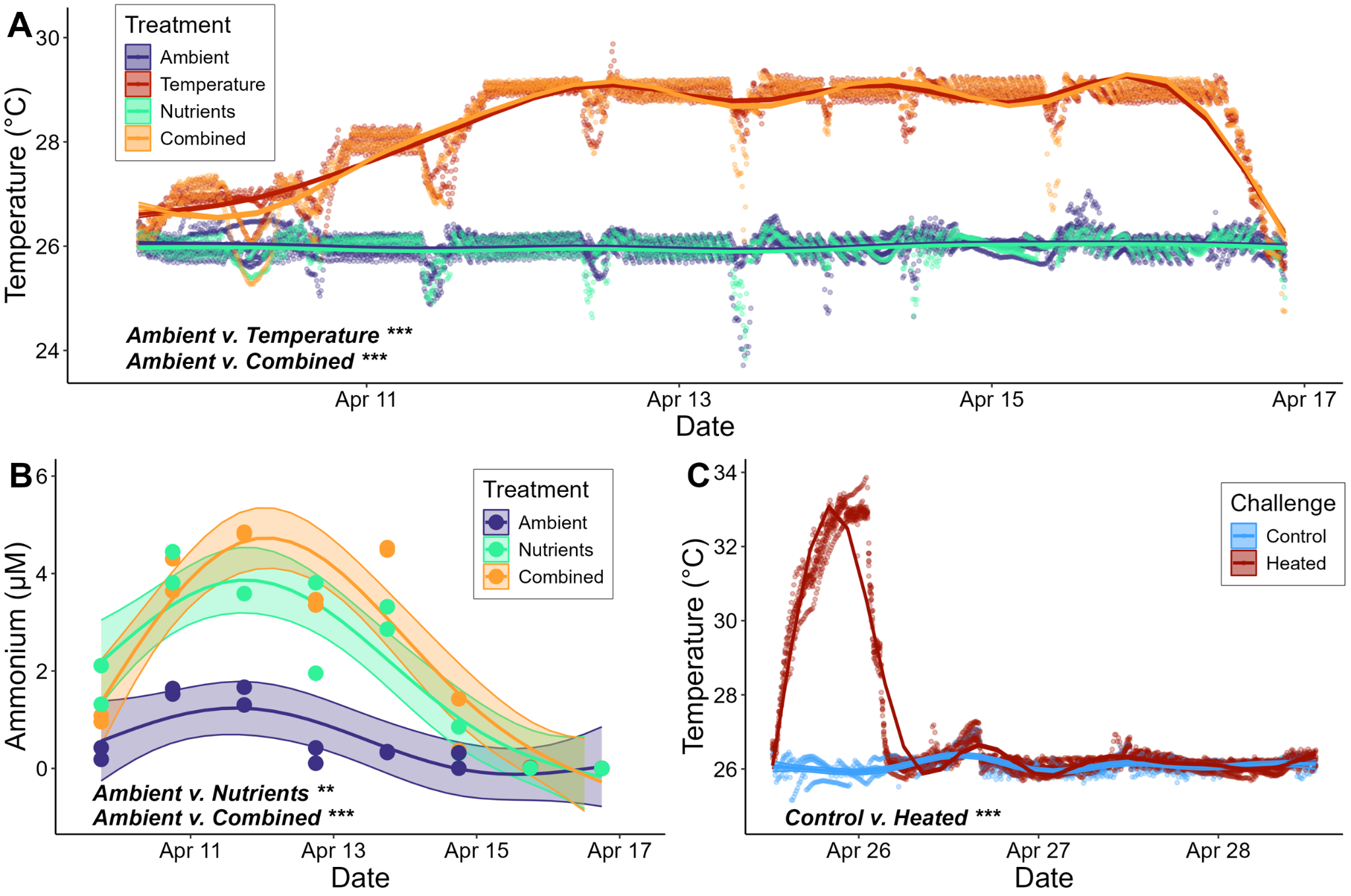

**Figure S3. Experimental conditions throughout priming and thermal challenge**. **A)** Temperature (℃) throughout the Priming phase. Points represent the water temperature (℃) measured every minute in each tank, averaged over 10-minute intervals. Temperature is compared between priming treatments (**Table S2A**). For panels **A-C**, lines represent the smooth terms from generalized additive models (GAM) and shaded regions show 95% confidence intervals. Points and lines are colored by treatment. Significant differences between treatments are noted with asterisks (*: p < 0.05, **: p < 0.01, ***: p < 0.001). **B)** Concentration of ammonium (NH_4_) throughout the Priming Phase. Points represent the concentration (μM) of ammonium measured each day (n=2 tanks sampled per treatment each day) and profiles over time are compared between priming treatments (**Table S2B**). **C)** Temperature (℃) throughout the Thermal Challenge. Points represent the 10 min averaged water temperature (℃) measured in each tank. Temperature is compared between Challenge treatments (**Table S2C**).

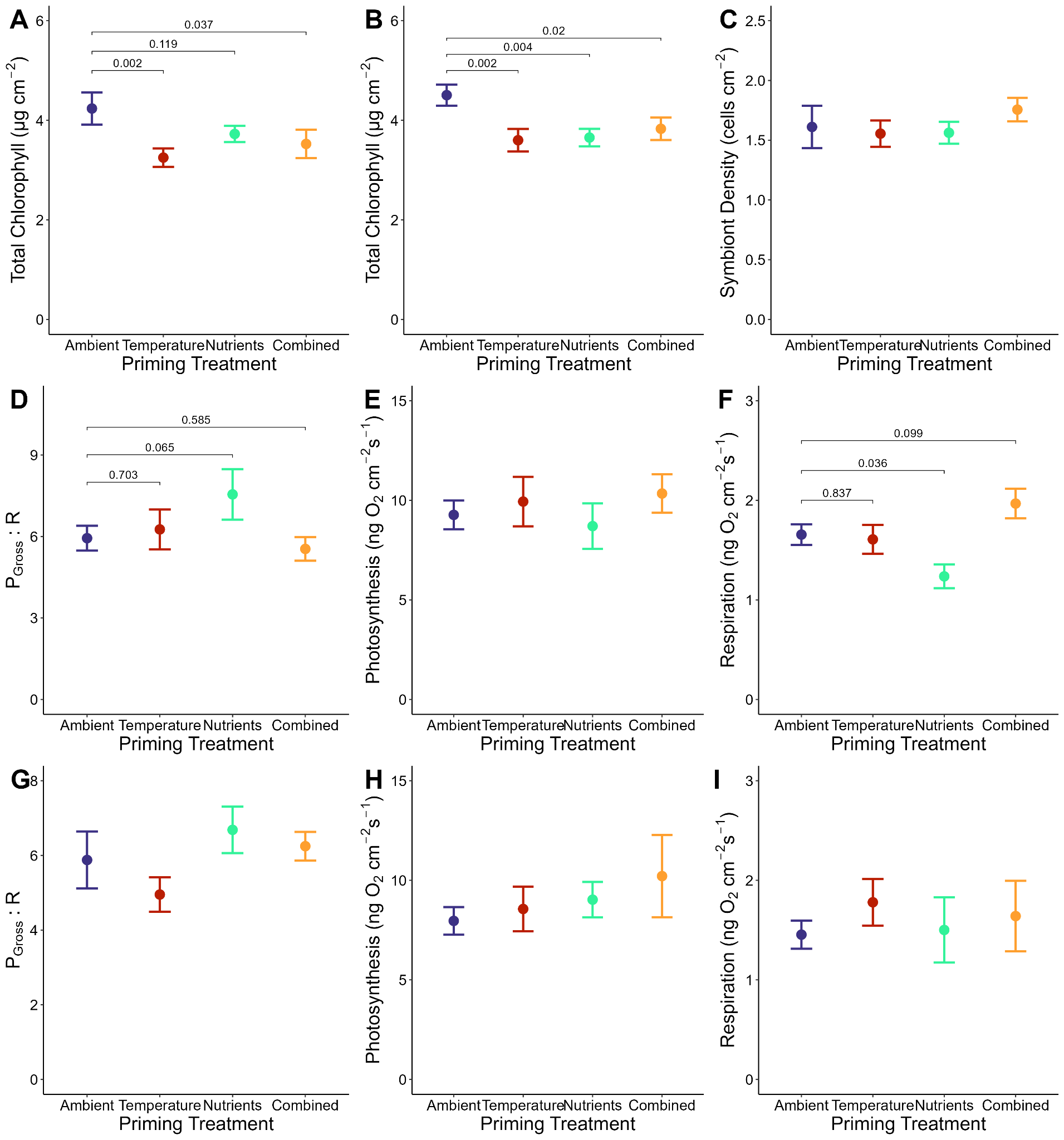

**Figure S4. Coral symbiosis and metabolism after priming and recovery.** Chlorophyll concentration (μg/cm^2^) for each treatment at the **A)** post-priming and **B)** recovery time points. **C)** Symbiont density (cells/cm^2^) post-priming. **D and G)** The ratio of gross photosynthesis to respiration (P_Gross_ : R), gross photosynthesis (ng O_2_/cm^2^/s), and **E and H)** respiration (ng O_2_/cm^2^/s) for each treatment at the mid-recovery **(D-F)** and full recovery **(G-I)** time points. Points represent averages, and error bars represent the standard error, each colored by priming treatment. For cases when the response metric was significantly influenced by the priming treatment, p-values comparing each primed treatment to the ambient control are shown for symbiosis and metabolism metrics **(Table S3 and Table S4, respectively)**.

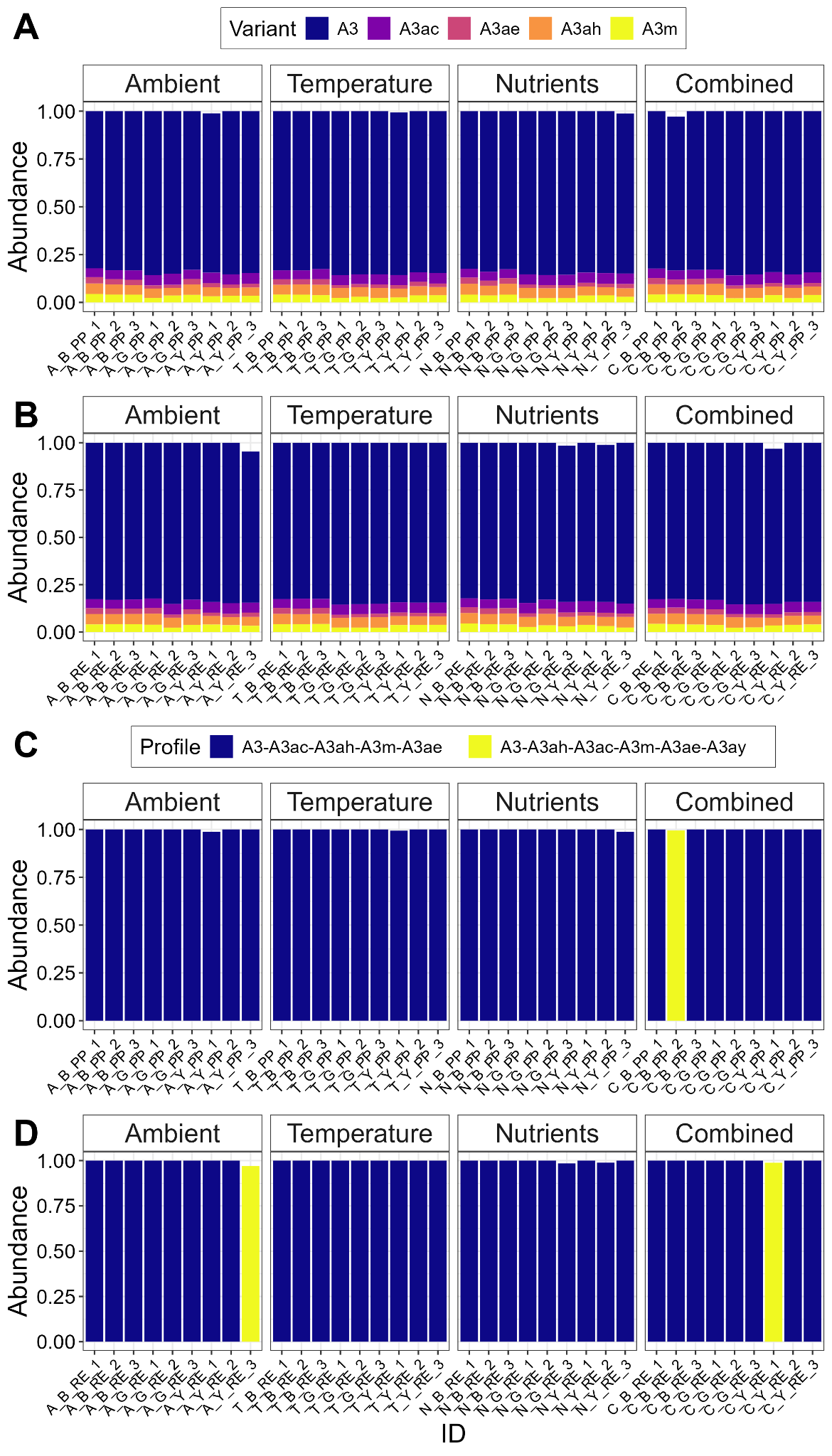

**Figure S5. Symbiont communities after stress priming.** Stacked barplots of the relative abundance of algal symbiont types at the level of **A-B)** sequence variants and **C-D)** profiles from ITS2 sequencing. Samples are organized by priming treatment (columns) and time point (**A** and **C**: post-priming, **B** and **D**: recovery). Results of community analysis are available in **Table S5**.

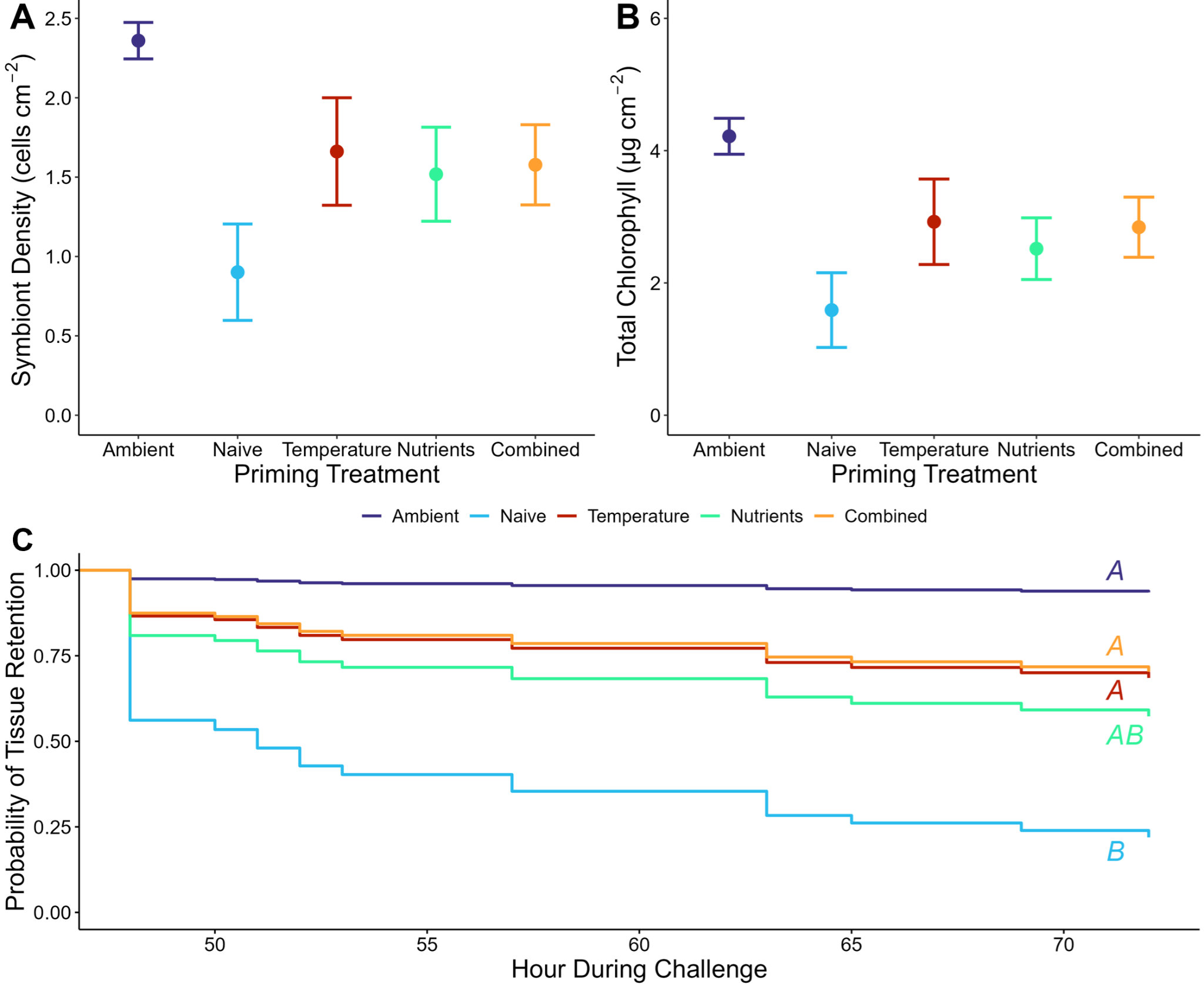

**Figure S6. Coral bleaching metrics during the thermal challenge.** **A)** Symbiont density (cells/cm^2^), **B)** chlorophyll concentration (μg/cm^2^) for each treatment at the post-challenge time point. Points represent averages and error bars represent the standard error, each colored by priming treatment. **C)** Survivorship curves (cox proportional hazard) of the probability of tissue retention throughout the thermal challenge. Curves are colored by priming treatment. Capital letters indicate significant pairwise differences in hazards between treatments (**Table S7**).

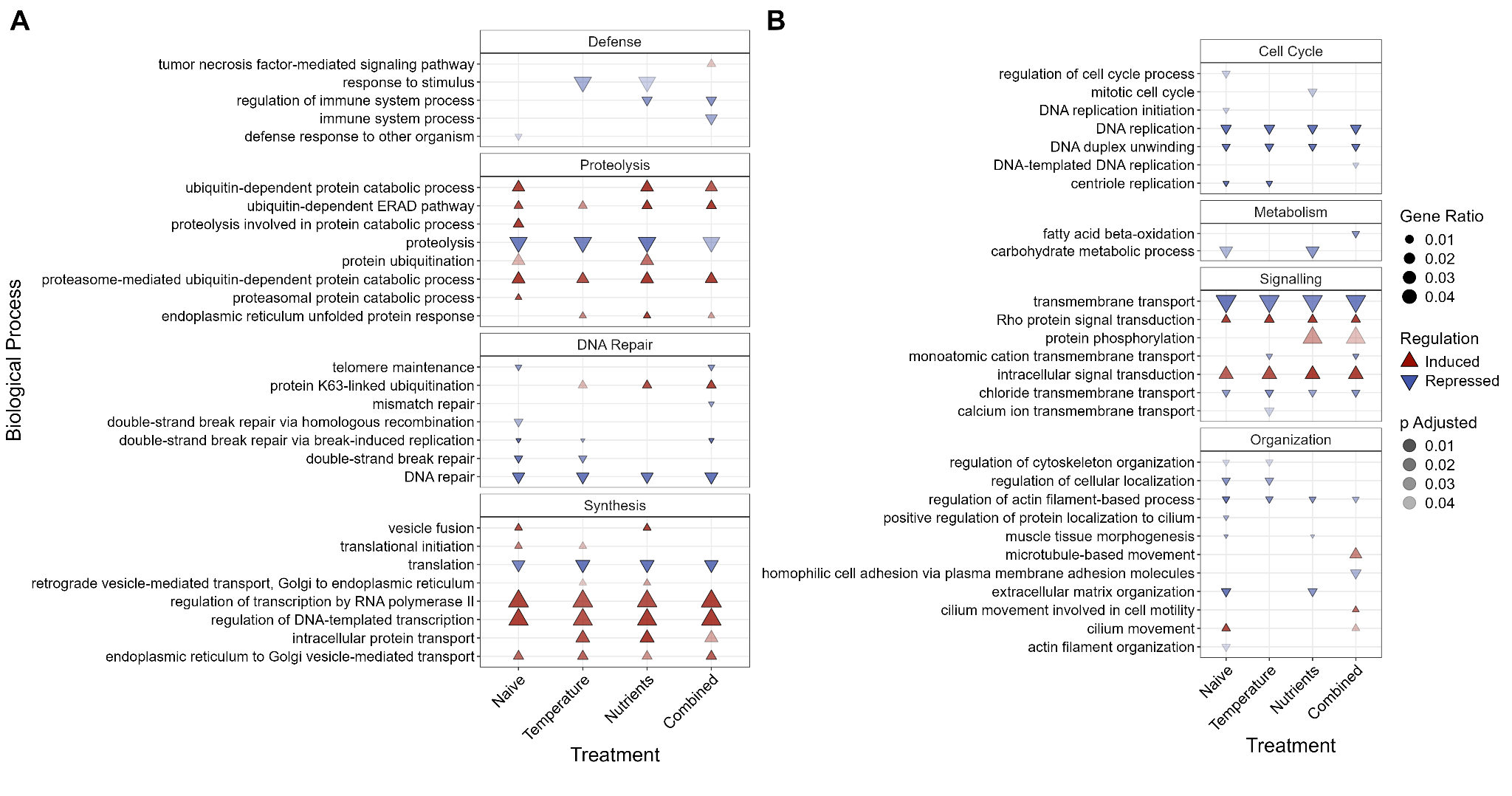

**Figure S7. Biological processes enriched at the end of the thermal challenge.** Gene ontology (GO) terms of biological processes enriched (**Table S8** ) in the heated treatments at the end of the thermal challenge. Induced terms are shown in upward red triangles and repressed terms are shown in downward blue triangles. The triangle size corresponds to the ratio of significant genes within the term and the transparency corresponds to the adjusted p value. GO terms are organized by similarity.

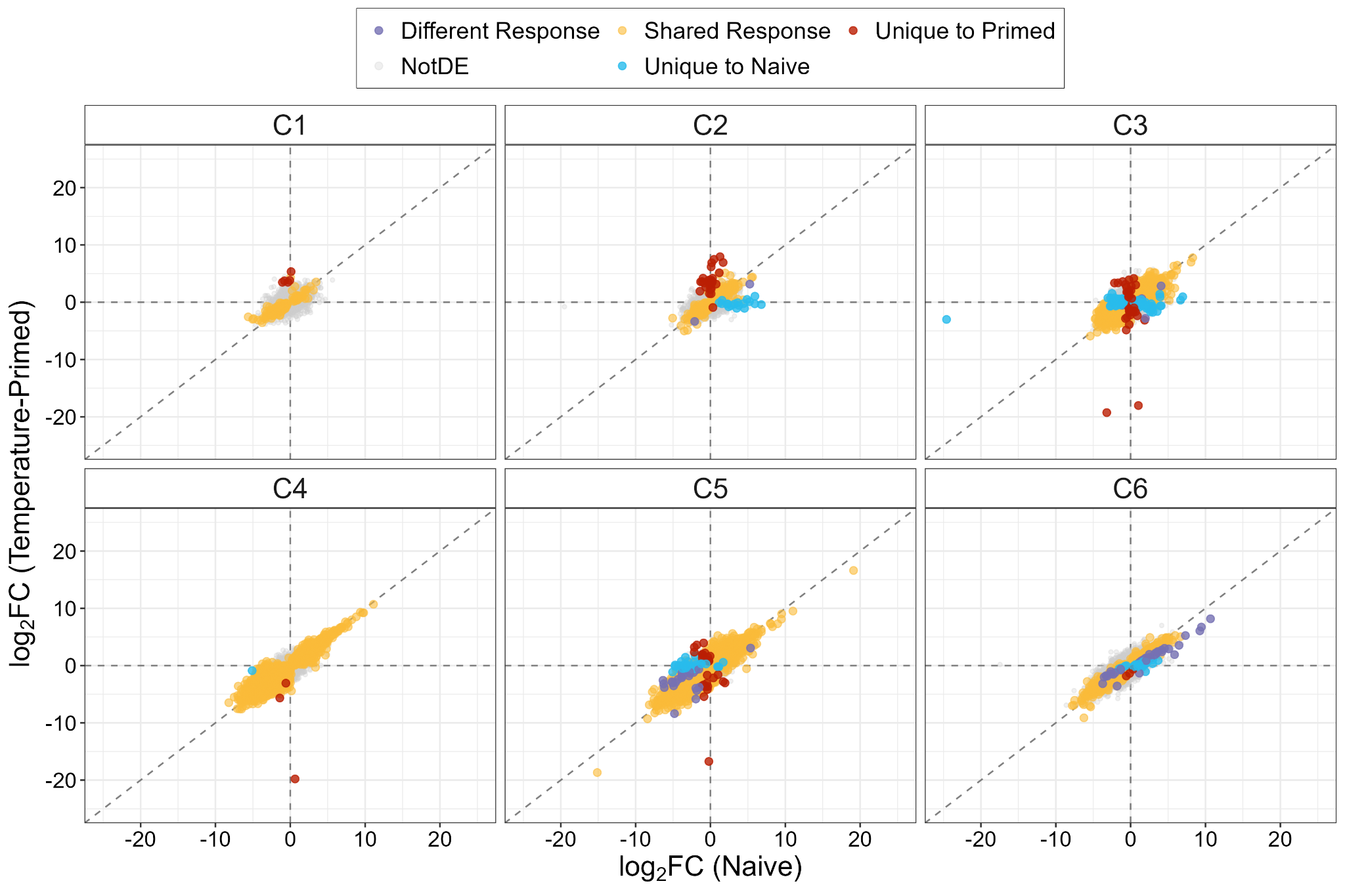

**Figure S8. Transcriptomic responses throughout acute heat stress.** Gene expression (log_2_ fold change [log_2_ FC]) of the naive corals versus the temperature-primed corals, relative to the ambient control corals. time points throughout the thermal challenge (C1-C6) are across panels. Points are colored by the uniqueness of the response between the naive and primed treatments.

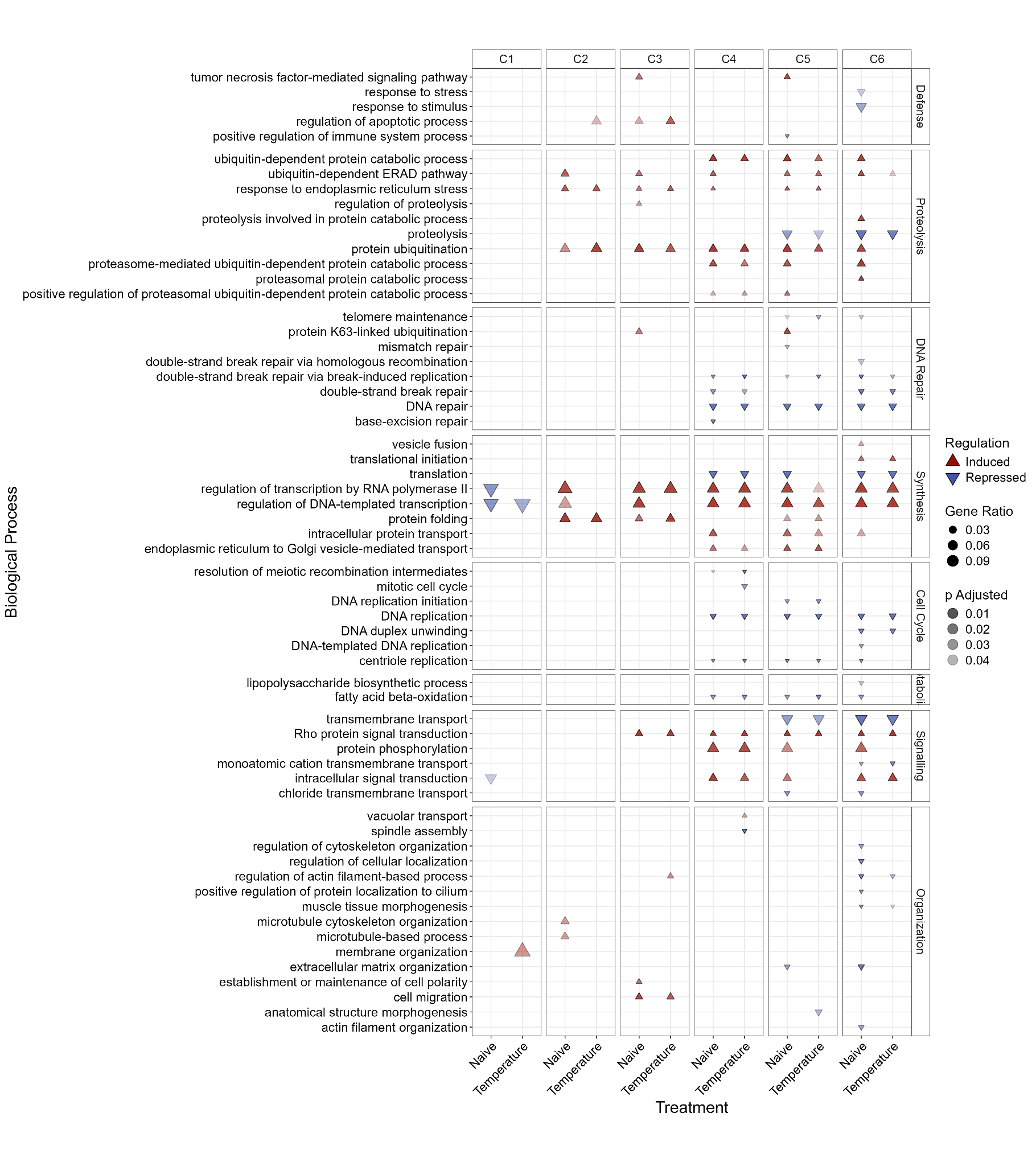

**Figure S9. Biological processes enriched throughout the thermal challenge.** Gene ontology (GO) terms of biological processes enriched (**Table S9** ) in the naive and temperature-primed treatments at each time point (C1-C6) throughout the thermal challenge. Induced terms are shown in upward red triangles and repressed terms are shown in downward blue triangles. The triangle size corresponds to the ratio of significant genes within the term and the transparency corresponds to the adjusted p value. GO terms are organized by similarity.

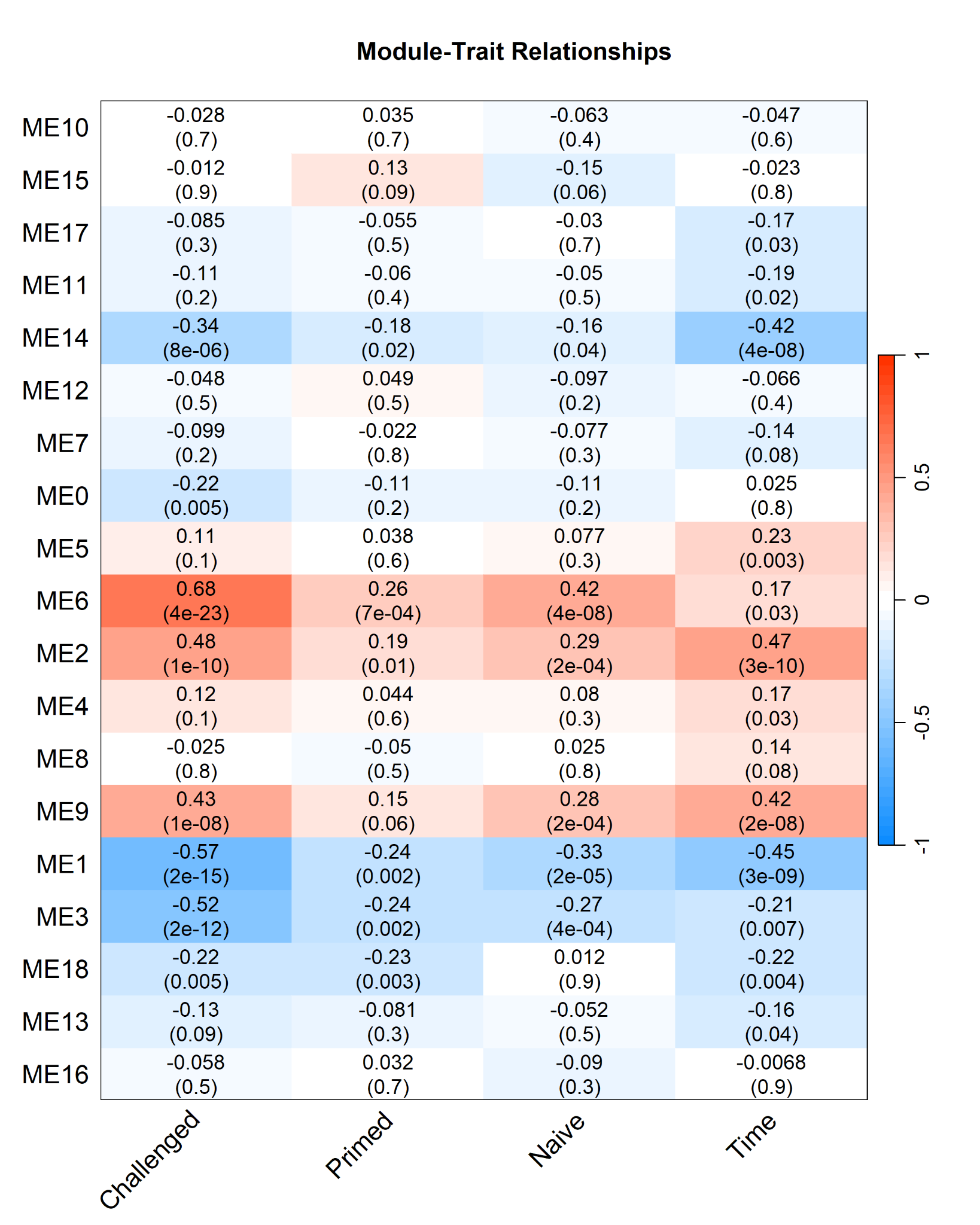

**Figure S10. WGCNA module and trait association heatmap.** Gene modules (y-axis) identified through weighted gene coexpression network analysis (WGCNA) and their correlations with four trait groups of interest: treatments heated during the thermal challenge (*i.e.,* both temperature-primed and naive), the temperature-primed or naive treatments, and time points throughout the acute heat exposure. Within each tile, correlations are shown, colored by positive (red) or negative (blue) correlations and p values are in parentheses.
